# Ecological and evolutionary dynamics of the oral microbiome across childhood

**DOI:** 10.64898/2026.02.13.705642

**Authors:** Fang Wang, Andrew J Holmes, Gina V Browne, Xuzhen He, Michelle R Bockmann, Kylie M Davis, Toby E Hughes, Christina J Adler

**Affiliations:** School of Medical Sciences, Faculty of Medicine and Health, The University of Sydney, Sydney, Australia; Charles Perkins Centre, The University of Sydney, Sydney, Australia; School of Life and Environmental Sciences, Faculty of Science, The University of Sydney, Sydney, Australia; Institute of Dental Research, WSLHD Research and Education Network, Westmead Hospital, Westmead, Sydney, Australia; School of Civil and Environmental Engineering, University of Technology Sydney, Sydney, Australia; School of Dentistry, Adelaide University, Adelaide, Australia; Faculty of Medicine and Health, The University of Sydney, Sydney, Australia

**Keywords:** human oral microbiome, microbial ecology, genetic diversity, metagenomics

## Abstract

Childhood represents a critical period for oral microbiome development, yet evolutionary trajectories and the relative roles of host and environment remain unclear. Using a large longitudinal metagenomic dataset of 920 samples from a twin cohort spanning the first decade of life, we characterised microbial shifts and population dynamics of key bacterial groups. Microbiome diversity was initially reduced and highly heterogeneous and became increasingly complex and convergent with age. Microbial community state was associated with developmental age, environment and in late childhood was surprisingly strongly associated with host genotype. Strain-level analyses revealed species-specific temporal patterns of genetic variation particularly within *Streptococcus*, reflecting adaptive responses to host and environmental pressures. *Fusobacterium* exhibited consistently high replication rates, indicating sustained growth dynamics. Phylogenetic reconstruction further revealed host and niche specific genomic diversification of Saccharibacteria lineages. These findings establish childhood as a decisive period of oral microbial evolution and highlight the role of host-microbiome and epithelial interactions in shaping community structure, providing guidance for oral management strategies that promote lifelong oral health.

## Main

Early life represents a critical window for oral microbiome acquisition, during which intense interactions among host factors, environmental exposures and microbial communities shape long-term microbial trajectories and oral ecosystem stability^1^. From birth through childhood, the oral microbiome assembles through a highly ordered process involving initial colonisation, microbial succession, and ecological maturation of the oral niches^2,3^. Early colonisers, particularly *Streptococcus* species, play a central role in this process by establishing foundational biofilm structures that facilitate subsequent community assembly and influence later disease risk^3,4^. Oral microbial homeostasis is essential not only for maintaining oral health but is increasingly linked to systemic conditions including metabolic, cardiovascular, and immune-mediated diseases^5–7^.

The dynamic development of the oral microbiome is shaped by the combined influences of host-regulated traits and environmental factors across early life. Host genetic factors may influence microbial colonisation through immune function, salivary composition, epithelial receptor expression, and oral anatomy^8,9^, whereas environmental influences including diet, oral hygiene practices, medication exposure, and lifestyle can begin even prenatally^10–12^. Disentangling the relative contributions of host genetics and environment across developmental stages is therefore critical in understanding oral microbiome development and informing oral health management strategies yet remains constrained due to limited longitudinal data and insufficient taxonomic and functional resolution.

Twin studies provide a powerful framework for inferring the relative contributions of additive genetic and environmental factors to human microbiome variation. In the gut, microbial taxa influenced by heritability which is defined as the proportion of variance attributable to host genetic differences have been associated with host genes involved in immune regulation and metabolism^13^. Similarly, oral microbiome studies using salivary samples have identified taxa influenced highly by host genetics beyond early childhood and reported associations with host genetic loci independent of cohabitation effects^14^. Relatively stable host genetic influence in oral microbiome diversity was reported in children aged 5-11 years alongside substantial temporal variability in heritability estimates for core individual oral taxa^15^. In contrast, environmental factors have also been identified as dominant drivers of oral microbial diversity, emphasising the role of age-related behaviours and individual oral hygiene practices in children aged 5-12 years^16^. These discrepancies likely reflect the dynamic nature of the oral microbiome, which varies with developmental stage, population characteristics, oral niche, and study design. While large-scale, longitudinal, metagenomic studies have advanced our understanding of gut microbiome development^17,18^, comparable studies of the oral microbiome remain scarce. Notably, previous twin studies of the paediatric oral microbiome have been limited by small sample sizes, cross-sectional designs, reliance on 16S rRNA profiling, and a lack of longitudinal and functional level resolution across key developmental windows hinder the power to capture the complex ecological and evolutionary dynamics of the oral microbiome^19–21^.

In this study, we addressed these gaps by leveraging a large longitudinal metagenomic dataset from a twin cohort spanning the first decade of life to characterise the evolutionary dynamics of the oral microbiome. Specifically, we aim to: 1) identify oral microbial shifts and key features linked with developmental stages; 2) quantify the relative contributions of host and environmental factors across childhood for both overall community structure and individual ecological features, including taxonomic and functional profiles; and 3) assess strain-level genetic variation influenced by temporal effect in key early colonisers and investigate previously under-characterised lineages at high resolution. Together, these analyses are essential for advancing our understanding of how microbial communities establish, adapt, and interact during early life as well as to guide efficient intervention strategies for oral and systemic health.

## Results

We retained 920 metagenomic samples after quality control, excluding those with excessive host contamination, abnormal read counts, or antibiotic use. These samples were selected from an initial 964 metagenomic sequencing libraries generated from oral biofilm samples collected from twin individuals (Methods). The samples were collected from 189 twin families from 4 Australia states at three different timepoints along their development (Fig. 1a, Supplementary Table 1), namely T1 (edentulous (no teeth), average age 6.8 ± 2.7 months, 266 samples), T2 (primary/deciduous/baby teeth only, 1.6 ± 0.4 years, 312 samples) and T3 (mixed dentition, 8.8 ± 1.2 years, 342 samples). A portion of these datasets had been utilized in our previous study with sequencing methods validated^22^. In total, metagenomic sequencing of DNA samples generated 8.67 trillion bp data and 57.8 billion reads for the 920 samples with an average sequencing depth of 31.4 million paired-end reads per sample (Supplementary Table 2).

**Fig. 1:**
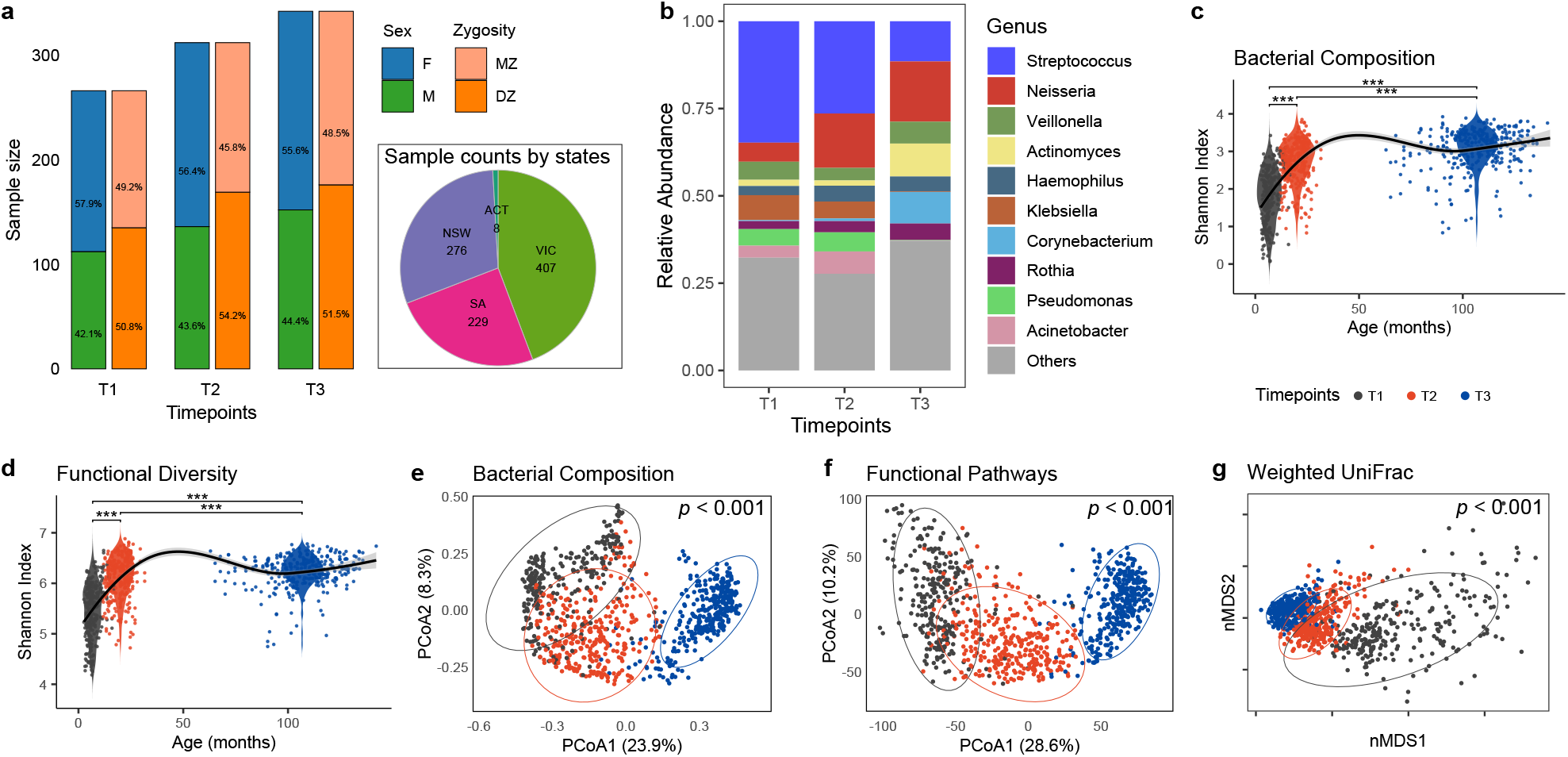
Longitudinal twin cohort dataset and overall divergence of oral microbiome across childhood. **a**, Sample sizes across the three timepoints in the cohort, stratified by sex and zygosity (T1 = 266, T2 = 312, T3 = 342. F = female, M = male, MZ = monozygotic, DZ = dizygotic). **b**, Relative abundance of the ten most abundant genera across the three timepoints. **c-d**, Alpha diversity (Shannon index) showing developmental shifts in bacterial community composition (**c**) and functional profiling based on species-stratified pathways after 0.01% filtering (**d**). Asterisks indicate significance from Tukey’s multiple comparisons test (\**p* ≤ 0.05, \*\**p* ≤ 0.01, \*\*\**p* ≤ 0.001). **e-g**, Beta diversity visualised by (**e**) PCoA of Bray-Curtis distances for bacterial composition, (**f**) PCoA of Euclidean distances for functional pathways with batch effect correction (Methods), and (**g**) non-metric multidimensional scaling (nMDS) of weighted UniFrac distances. Statistical differences across timepoints were assessed using PERMANOVA, adjusting for sex and repeated measures within families. Ellipses represent 95% confidence intervals.

We leveraged both read-based analysis and assembled Metagenome-Assembled Genomes (MAGs) to investigate different aspects of the development of the oral microbiome across three timepoints (Fig. 2). Reads-based analyses provided broad insights into microbial community patterns, whereas MAGs offer genome-level insights for particular bacteria with high resolution.

**Fig. 2:**
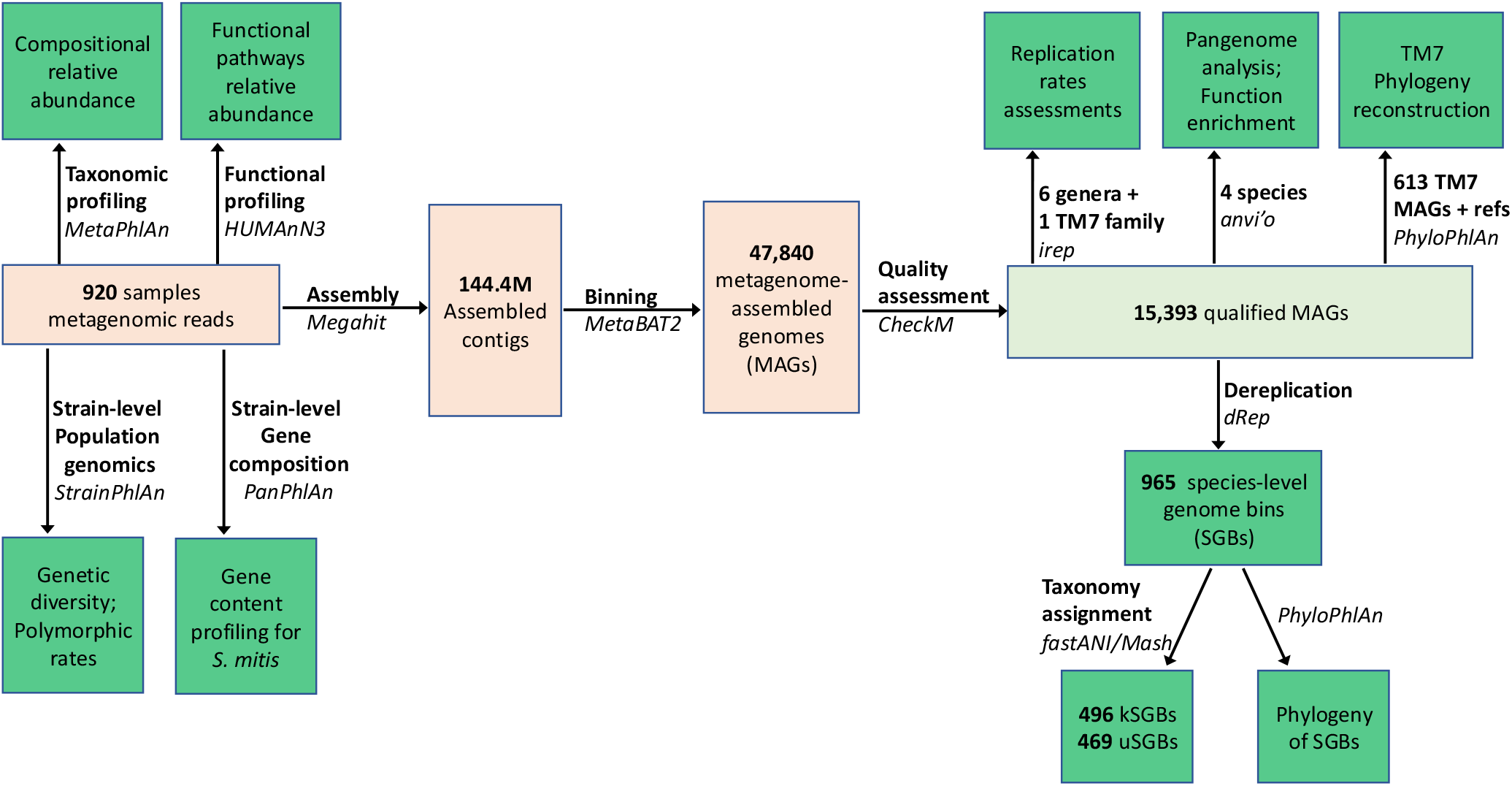
Overview of data processing and analytical workflow.

### Reads-based analysis

#### Divergence in composition and functional pathways in the children oral microbiome

Read-based relative abundance analysis revealed distinct temporal shifts in both the ecological composition and functional capacity (metabolic pathways inferred from microbial gene contents) across the first decade of life. At six months of age (T1), *Streptococcus* dominated the community with 34.8% of total sequence reads. This decreased to 26.4% by two years of age (T2) and declined to 11.5% by ages five to eight years (T3), accompanied by an emergence of a more diverse microbial assemblage, including *Neisseria* (17.3%), *Actinomyces* (9.4%), and *Corynebacterium* (9.0%) (Fig. 1b). Predominant functional capabilities of overall community remained consistent across T1, T2, and T3.

Our data showed the sampled habitats supported an oral microbial community of increasing complexity over time. The overall alpha diversity increased significantly at both the ecological level (bacterial composition, mean Shannon index: T1-1.80, T2-2.72, T3-3.07) and the functional level (species-stratified metabolic pathways, T1-5.45, T2-6.12, T3-6.23) (Fig. 1c-d). Timepoint had a strong and statistically significant effect on alpha diversity based on linear mixed-effects models that adjusted for sex and accounted for within-family correlations (analysis of variance (ANOVA), *p* < 0.0001 for both). All pairwise differences between timepoints remained statistically significant after Tukey-adjusted comparisons (*p* < 0.0001 for all contrasts). No significant effect of sex on alpha diversity was observed.

Beta diversity analysis revealed significant differences in oral community structure across timepoints. Principal Coordinates Analysis (PCoA) and Permutational Multivariate Analysis of Variance (PERMANOVA) demonstrated strong and significant separation by timepoint for both microbial composition and functional potential (Fig. 1e-f; R^2^=0.23, 0.29; p < 0.001 for both), adjusting for sex and family effects and accounting for repeated measures within individuals. We employed the weighted-unifrac distance metric to investigate the degree of ecological divergence of microbial communities across childhood, considering both the relative abundances of each species and phylogenetic relationship among them. A distinct segregation of samples was observed by timepoint (Fig. 1g) with significance (PERMANOVA: R^2^=0.34, p < 0.001). Multivariate dispersion also differed significantly among timepoints (ANOVA, permutest, all p < 0.001), with T1 showing greater variability and more outliers compared to T2 and T3. This pattern was most pronounced for UniFrac distances (Average distance to group centroid: T1-0.39, T2-0.28, T3-0.21), indicating increasing phylogenetic convergence of the oral microbiome over time. Overall, the analysis suggests that the oral microbiome in infancy is characterised by lower diversity and higher inter-individual variability. As children grow, their oral microbiomes become more diverse within individuals (increased alpha diversity) yet more similar across individuals (reduced beta diversity), reflecting the maturation and stabilisation of the microbial community. The strong concordance between species-level taxonomic shifts and changes in functional pathways supports the robustness of the dataset for downstream ecological and genetic inference.

#### Microbial species and functional signatures linked with children age

To evaluate whether oral community assembly outcomes at different ages are predictable, we constructed classification models using Random Forest algorithms based on species or functional (metabolic pathway level) datasets. Six datasets were analysed, including full feature sets and subsets comprising only differentially abundant features (Methods).

Both taxonomic and functional profiles discriminated temporal variation effectively with high classification accuracy (Fig. 3a, Supplementary Table 3). Species-level and species-stratified pathway profiles achieved the highest performance (>95% accuracy), demonstrating strong temporal signals in both microbial community structure and functional potential. Overall, the OOB (out of bag) error rates were below 0.05, with highest accuracy in predicting T3 and lowest in T1 (Fig. 3b, Extended Data Fig. 1a), potentially due to the consistent and stable microbial profile in T3 shown above.

**Fig. 3:**
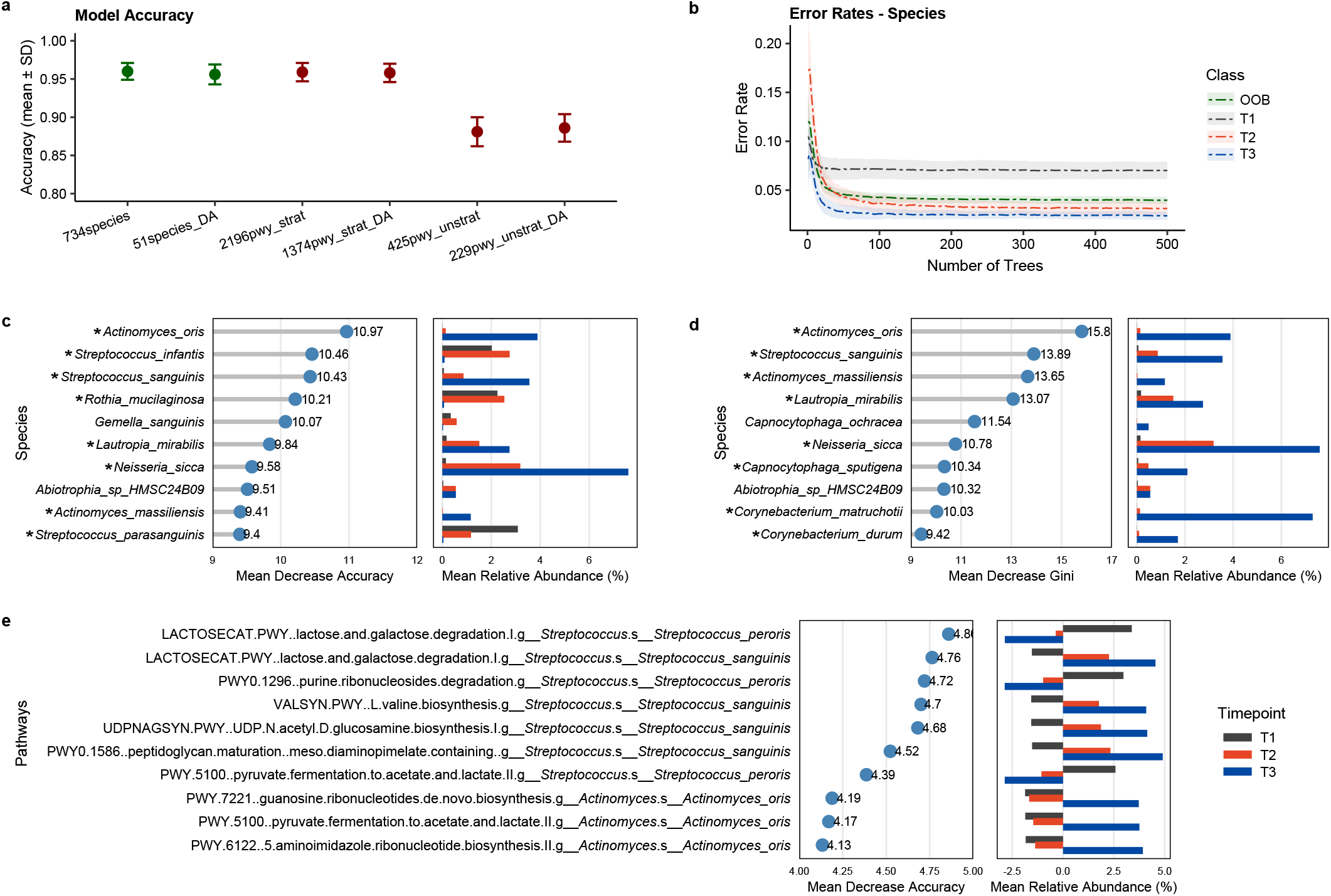
Classification models and key features associated with timepoints. **a**, Accuracy of random forest models based on six sets of species and pathways abundance profiles. DA datasets referred to features identified as differentially abundant by MaAsLin2. **b**, Error rate plot showing the mean and standard deviation across 100 random forest runs, based on abundance profiles of 734 species (trained on 75% of the samples, tested on the remaining 25%; split stratified by timepoint; 500 trees per model). OOB: out-of-bag error estimate from internal cross-validation within the training set. **c**, Top ten species ranked by variable importance in the random forest model based on abundance profiles of 734 species assessed by Mean Decrease Accuracy (reduction in classification accuracy when each feature is permuted across samples). Asterisks denote species identified as differentially abundant among timepoints by MaAsLin2. Left panel: Mean Decrease Accuracy for each species; right panel: mean relative abundance across samples per timepoint. **d**, Top ten species ranked by Mean Decrease Gini which reflects the total reduction in Gini impurity contributed by each feature across all decision trees, based on abundance profiles of 734 species. **e**, Top ten species-stratified pathways ranked by Mean Decrease Accuracy.

The life-history related species-stratified functional traits outperform shared functional traits. This highlights that successional outcomes in the oral microbiome are driven primarily by biological processes that are more strongly dependent on organism-level activities and adaptations rather than shared community-level functions. Additionally, the differentially abundant feature subsets retained accuracy comparable to the full feature profiles, indicating that early-life microbiome succession is driven by a focused group of key taxa and pathways, whereas many other features contribute redundant or minimal predictive information. Consistently, the top predictive variables identified from the full species dataset were also among the differentially abundant taxa, including *Actinomyces oris, S. sanguinis, Actinomyces massiliensis*, and *Lautropia mirabilis* (Fig. 3c-d) with most of them enriched at T3.

#### Shifting key determinants of oral microbiome diversity across childhood

The large twin study design enabled the estimation of driving forces of the oral microbiome diversity at each timepoint. Using a univariate ACE structural equation modelling^23,24^, we were able to quantify the relative contributions of additive genetic effects or heritability (A), shared environmental effects (C), and unique environmental effects (E) to variations in the oral microbiome. This approach assumes that monozygotic (MZ) twins share around 100% of their genes, whereas dizygotic (DZ) twins share on average 50%, and that shared environmental exposures influence both twin types to a similar extent. Thus, greater within-pair correlation in MZ twins compared with DZ twins is attributed to genetic influences, whereas similarities in MZ and DZ correlations reflect shared environmental effects. Residual differences within twin pairs are attributed to unique environmental effects plus measurement error.

We tested for alpha diversity of the overall community at each timepoint. Both compositional and functional diversity displayed similar patterns with environmental factors dominated at early timepoints (T1, T2), whereas heritability became primary drivers at T3 (Fig. 4a). For compositional diversity, heritability was negligible at T1 and T2, with shared environmental effects explaining 63% (T1) and 52% (T2) of the variance. By T3, the estimated heritability increased markedly (50%) while shared environmental influences dropped substantially. Functional diversity followed the same trend with minimal heritability at T1 (A = 0%) and T2 (A = 1.5%) and shared environment dominating (66% and 65%). By T3, heritability rose sharply to 55% while shared environmental influences dropped to 5%. Unique environmental influences remained relatively stable across childhood for both compositional and functional diversity. These findings indicate that heritability increasingly modulates the oral microbiome as children age, overtaking the influence of shared environmental factors. This is potentially associated with the maturation of the oral epithelium and salivary environment and immune system as children age which introduce host-specific binding sites, immune pressures, and biochemical constraints that increasingly shape microbial colonisation and persistence. Together, these host-driven factors promote the establishment of stable, individual-specific oral biofilm communities and regulate the long-term incorporation of newly encountered microbes alongside ongoing environmental exposure.

**Fig. 4:**
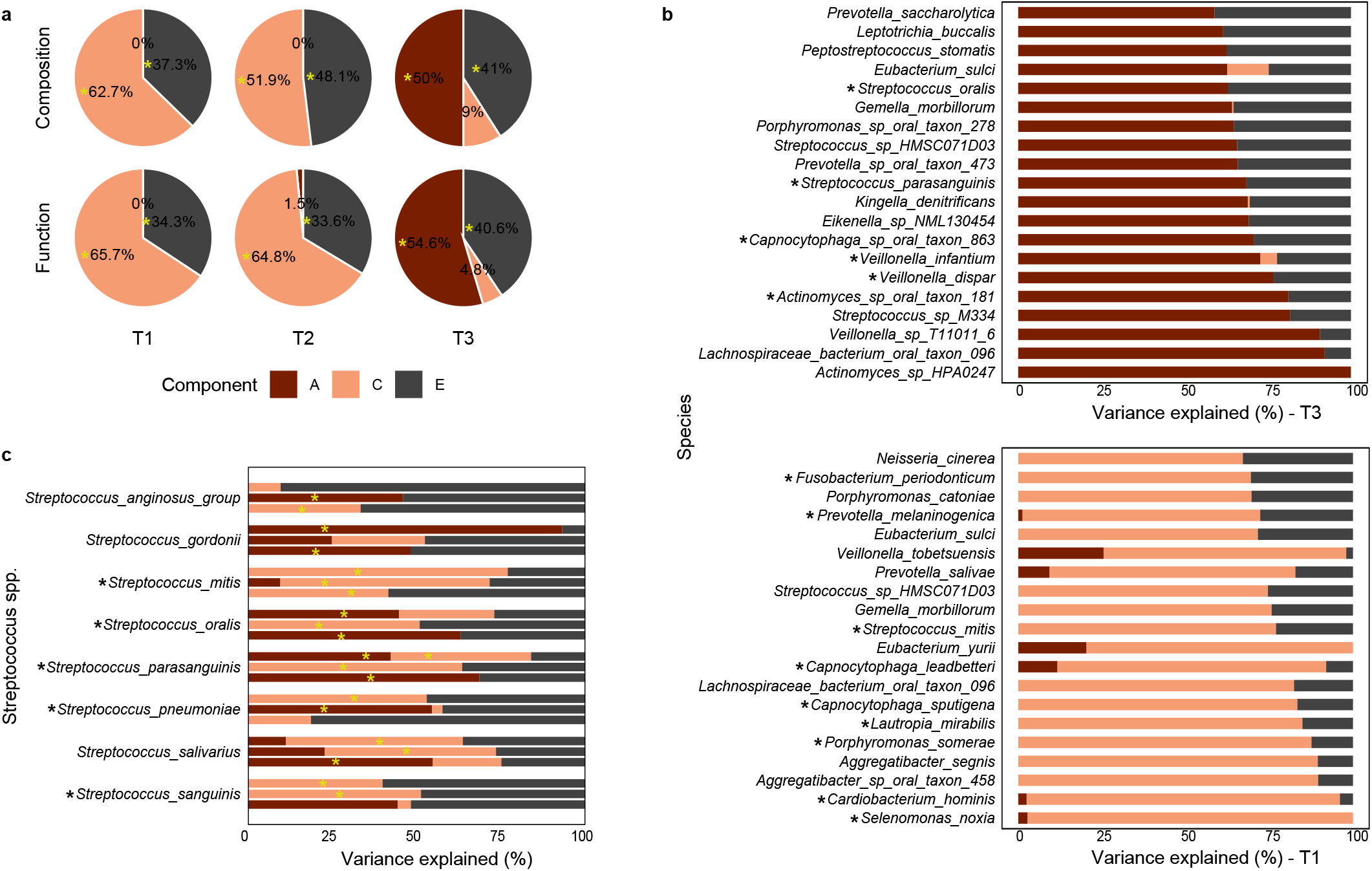
Host genetic and environmental contributions to microbiome dynamics. **a**, Partitioning of host genetic and environmental contributions to overall microbiome alpha diversity, shown separately for community composition (top row) and predicted function (bottom row) at each timepoint. Each pie chart is rescaled to 100% to facilitate comparison across timepoints and components. A = additive genetic, C = shared environmental, E = unique environmental. Yellow asterisks indicate significance of inclusion necessity of each component evaluated by model comparison using likelihood-ratio tests. **b**, Partitioning of host genetic and environmental contributions for species most strongly influenced by host genetics (A) at T3 (top) and by shared environmental factors (C) at T1 (bottom). Species are ordered vertically according to the proportion of additive genetic (A) or shared environmental (C) contributions. **c**, Host genetic and environmental contributions to *Streptococcus* species across three timepoints. Asterisks beside the species indicate species identified as differentially abundant among timepoints. Yellow asterisks indicate significance of necessity for inclusion of a component in the model. Species are ordered alphabetically with T1-T3 shown from top to bottom.

#### Microbial species and function signatures influenced by host genetics and environmental factors

Using ACE modelling, we further assessed genetic and environmental contributions to individual microbial species and functional pathways across childhood to identify features strongly influenced by host genetics at T3 and environmental factors at T1.

At T3, several species exhibited strong host genetic influence over their persistence, including *Actinomyces* sp. HPA0247, *Lachnospiraceae* bacterium, and *Veillonella* sp. T11011-6. Many other species from abundant genera such as *Streptococcus, Veillonella*, and *Actinomyces* were also strongly influenced by host genetics. While primarily commensal, some taxa (*Porphyromonas* sp. PRA taxon 278, *Eikenella* sp. NML130454, *Peptostreptococcus stomatis*) have been implicated in periodontitis under dysbiotic conditions (Fig. 4b). At the functional level, host genetics contributed more than 85% of the variance in core microbial functions including L-tyrosine biosynthesis (PWY-6630), polyamine biosynthesis II (POLYAMINSYN3.PWY), and the S-adenosyl-L-methionine cycle (PWY-6151) (Extended Data Fig. 2). In contrast, at T1, species such as *Selenomonas noxia, Cardiobacterium hominis, Aggregatibacter* spp., and *Porphyromonas somerae* were predominantly shaped by shared environmental factors, reflecting sensitivity to household-level exposures. *L. mirabilis*, also identified as an important predictive species, was influenced highly by shared environmental influence and negligible by host genetics (Fig. 4b). Pathways including isoprene biosynthesis II (PWY-7391), pyrimidine deoxyribonucleotide biosynthesis I (PWY-7184), and pyrimidine deoxyribonucleotides de novo biosynthesis II (PWY0-166) were largely influenced by shared environment (>70%) at T1.

Temporal species-specific patterns were also evident in early colonisers. *S. mitis* and *S. sanguinis* were mainly shaped by environmental factors at early timepoints, with *S. sanguinis* later becoming more influenced by heritability. In contrast, *S. gordonii, S. oralis*, and *S. parasanguinis* exhibited strong impact by heritability at T1 (Fig. 4c). In contrast to previous reports suggesting that Streptococcus species are predominantly influenced by environmental factors during mid-childhood^15^, we observed that several Streptococcus taxa displayed increased heritability at T3, including *S. parasanguinis, S. oralis*, and *S. salivarius*. Among *Actinomyces, A. johnsonii* variation was entirely attributable to environmental factors throughout childhood, whereas *Fusobacterium nucleatum* was strongly influenced by heritability especially at early timepoints (Extended Data Fig. 2a).

Differential responses of oral species to host and environmental factors over time reflect species-specific niche adaptation, life-history strategies, and interaction networks, combined with developmental changes in the host oral environment that increasingly impose deterministic ecological filtering during microbiome succession.

#### Within-species genetic variation of major oral species

Taking advantage of the large-scale metagenomic sequencing data, we explored how oral microbial populations evolve during childhood by conducting strain-level analyses to characterise within-species genetic variation and its relationship with dentition development. We examined 44 bacterial species (see methods) on their population genetic features. For each species, we quantified intraspecific variability using (1) genetic diversity, assessed from pairwise genetic distances among consensus genomes, and (2) polymorphism rates, measured as the frequency of polymorphic sites across species-specific marker genes.

Pairwise genetic distances revealed strong species-specific variation in strain-level diversity (Extended Data Fig. 3). Species, timepoint, and their interaction were all significant (two-way ANOVA, p < 2 × 10^−16^). When each species was examined individually, most of them exhibited significant temporal shifts in diversity (ANOVA, FDR-adjusted), except *Capnocytophaga ochracea* and *Serratia liquefaciens* which remained stable. These findings indicate distinct, species-specific trajectories of strain-level diversity with temporal signal. Genus-level analysis of *Neisseria* (8 species), *Streptococcus* (13 species), and *Veillonella* (4 species) showed that genus, timepoint, and their interactions significantly influenced diversity (two-way ANOVA, p < 2 × 10^−16^), with *Neisseria* exhibiting the highest within-genus diversity. When distances were averaged per species and timepoint (Supplementary Table 4), only species identity remained significant (explaining 89% of variance), whereas timepoint alone was not significant (ANOVA based on linear mixed model: F = 2.20, p = 0.12), suggesting that interspecies differences dominate overall variation, with no consistent global age-related trend once species identity is considered.

We next quantified polymorphic site frequencies as a finer measure of within-population genetic variation for each species^25^. Polymorphic rates defined as the median percentage of such sites across all marker genes for each species, ranged from 0.16%-6.87% (Fig. 5), with significant effects of species identity and timepoints (two-way ANOVA, p < 2 × 10^−16^). Kruskal–Wallis and Dunn’s post hoc tests (FDR < 0.05) confirmed temporal variation in most species (Supplementary Table 5). Notably, several *Streptococcus* species (*S. mitis, S. infantis, S. oralis, S. peroris*, and *Streptococcus mutans*) exhibited consistently high polymorphic rates across childhood, indicating elevated strain-level genomic diversity. This likely reflects strong niche adaptation arising from interactions with concurrent species and the host environments^26^.

**Fig. 5:**
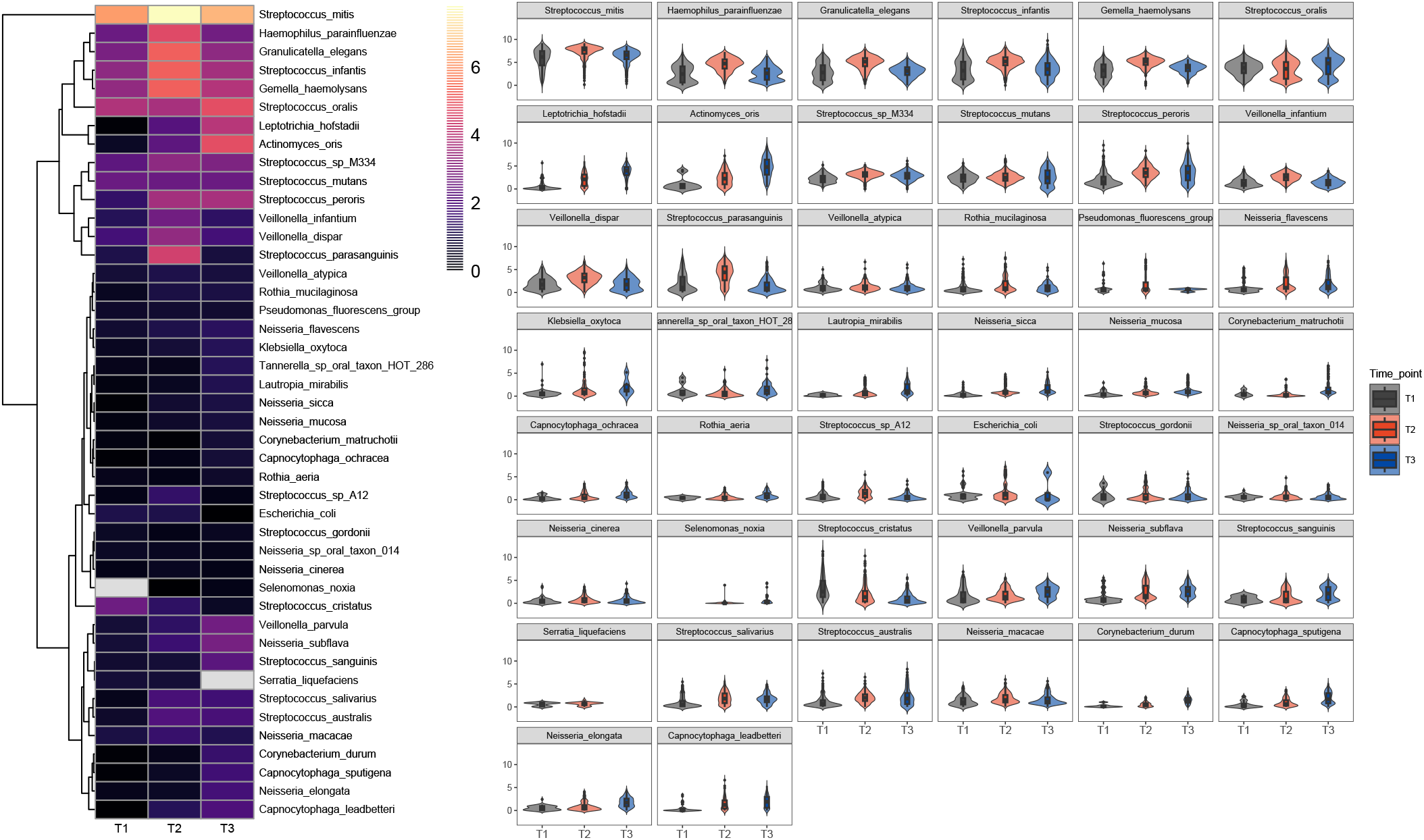
Polymorphic rate dynamics of oral species across timepoints. Polymorphic rates of 44 species across developmental timepoints. Left panel: heatmap of median polymorphic rates per species, clustered using Euclidean distance and complete linkage. Colours represent relative rates, with lighter shades indicating higher values. Blank cells denote absence of data for the corresponding species and timepoint; right panel: violin–box plots of polymorphic rates by timepoint for each species in a fixed y-axis scale. Violin shapes show the distribution of rates and embedded box plots indicate the median and 25^th^, 75^th^ range, with whiskers extending to 1.5 × IQR (Interquartile Range).

#### Temporal strain-level variation in *Streptococcus mitis* across childhood

We next examined strain-level variation in the prevalent oral species *S. mitis*. Analyses of genetic diversity, phylogeny, and gene-content profiles revealed clear temporal structuring across the three timepoints, supporting its adaptive role in early niche establishment. (Fig. 6). Phylogeny analysis using multiple sequence alignments across samples showed T3 strains largely occupying separate clades from strains of earlier timepoints (Fig. 6a), consistent with Principal Coordinate Analysis from distance metrics, where T3 samples form a distinct cluster from T1 and T2 (Fig. 6b). Genetic diversity differed significantly across timepoints (PERMANOVA, p = 0.001). Multivariate dispersion also varied significantly (betadisper, p = 0.001) indicating differences in within-group variability. One-way ANOSIM further supported a modest but significant temporal effect on strain-level genetic variation (R = 0.15, p = 0.001).

**Fig. 6:**
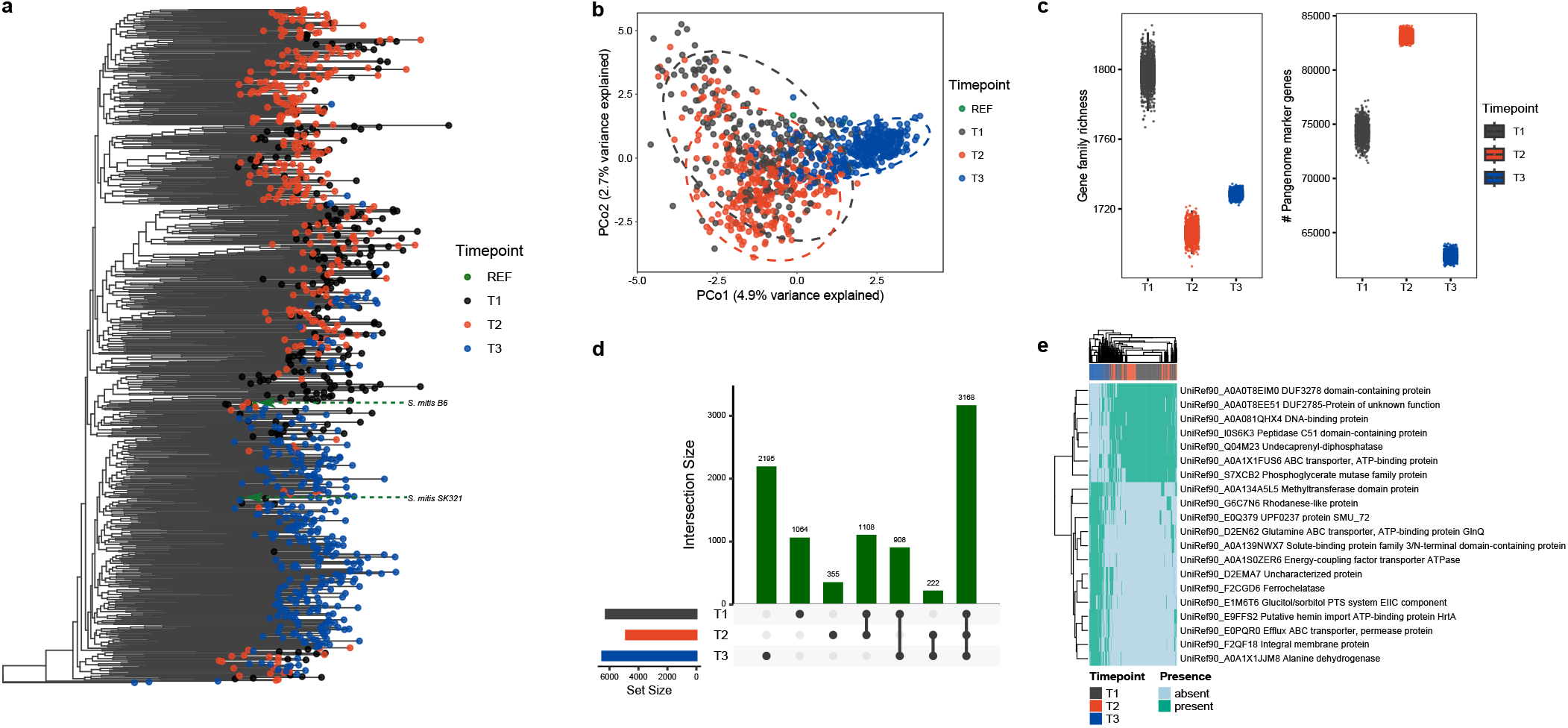
Genetic variation of *Streptococcus mitis* across childhood. **a**, RAxML phylogenetic tree of *S. mitis* strains, with tips coloured by sampling timepoint, illustrating temporal clustering. The number of strains included per timepoint was T1 = 244, T2 = 308, and T3 = 336. Two reference genomes were included for context and are indicated with green arrows: *S. mitis* B6 (GCF_000027165.1) and *S. mitis* SK321 (GCF_000148505.1). **b**, Principal Coordinates Analysis (PCoA) of pairwise genetic distances calculated from StrainPhlAn multiple-sequence alignments, showing clear separation of *S. mitis* populations by timepoint. **c**, Boxplots with overlaid jittered points showing (left) gene-family richness (number of UniRef90 clusters per sample) and (right) total pangenome marker genes detected per sample. Fill and point colour denote timepoint. Boxplots are based on balanced subsampling (n = 60 samples per timepoint, 1000 iterations) to account for unequal sample sizes, and show the distribution of richness across subsampling iterations. **d**, Upset plot of gene-family distributions across timepoints based on balanced subsampling (n = 60 samples per timepoint, 1000 iterations). Bottom horizontal bars (“set size”) represent the median values of total number of distinct gene families detected in at least one sample from each timepoint across iterations; top vertical bars (“intersection size”) show the number of gene families that are either unique to or shared between specific combinations of timepoints, denoted by the connected dots below. **e**, Heatmap of the top 20 *S. mitis* gene families that varied significantly across timepoints. Rows represent gene families (labelled with UniRef90 ID and functional annotation), and columns represent individual samples ordered by hierarchical clustering of gene presence/absence profiles.

We next performed strain-specific gene-content profiling of *S. mitis* using a pangenome-based approach to assess whether dominant strain composition shifted over time. Pronounced temporal differences were observed in the *S. mitis* accessory genome (Fig. 6c-d). On a per-sample basis, gene-family richness peaked at T1 and was lowest at T2, despite T2 samples exhibiting the highest pangenome marker gene counts. This pattern suggested that T2 strains concentrated within a smaller set of shared gene families. By contrast, although individual T3 samples contained fewer marker genes on average, the collective T3 repertoire harboured the largest number of unique accessory gene families. These results suggested increasing lineage diversification of the *S. mitis* population at T3, potentially reflecting adaptive responses to the more complex and dynamic oral environment during later stages of children oral development^27^.

Of the 9,727 profiled gene families, 1,132 displayed significant differences in presence across the three sampling timepoints (logistic regression, FDR-adjusted; see Methods). The top 20 most differentially present families revealed enrichment of early-colonisation accessory functions in T1–T2 samples, whereas T3 samples displayed enhanced metabolic flexibility and stress response capabilities in *S. mitis* (Fig. 6e).

### MAGs-based analysis

#### Reconstruction of microbial genomes from oral metagenomic sequencing data

From the 920 oral metagenomic samples, 144.4 million contigs were assembled from the targeted reads (mean 156,324 per sample; mean length 1,360 bp; N50 = 4,386 bp). This resulted in 47,840 reconstructed MAGs, of which 15,393 met quality criteria^28^ with 5,396 classified as high-quality (HQ) and 9,997 as medium-quality (MQ) (Methods; Fig. 2; Supplementary Table 6). All downstream analyses were based on the 15,393 HQ and MQ MAGs. Dereplication of these qualified MAGs with dRep^29^ resulted in 965 representative species-level genome bins (SGBs) spanning 13 phyla, with Pseudomonadota, Bacillota, Actinomycetota, and Bacteroidota being the most prevalent (Fig. 7, Supplementary Table 7). Among the three timepoints, T3 yielded the highest number of MAGs per sample. Taxonomic assignment identified 496 known SGBs (kSGBs), while the remaining 469 unclassified SGBs (uSGBs) had no matches in reference databases, most of which belonged to Pseudomonadota (31.8%) and Bacillota (27.5%). Phylogenetic reconstruction largely mirrored phylum-level taxonomy and aided in classification, although a small subset of MAGs mainly MQ and family-level genome bins showed deviations, reflecting potential novel lineages or incomplete genomes^30^.

**Fig. 7:**
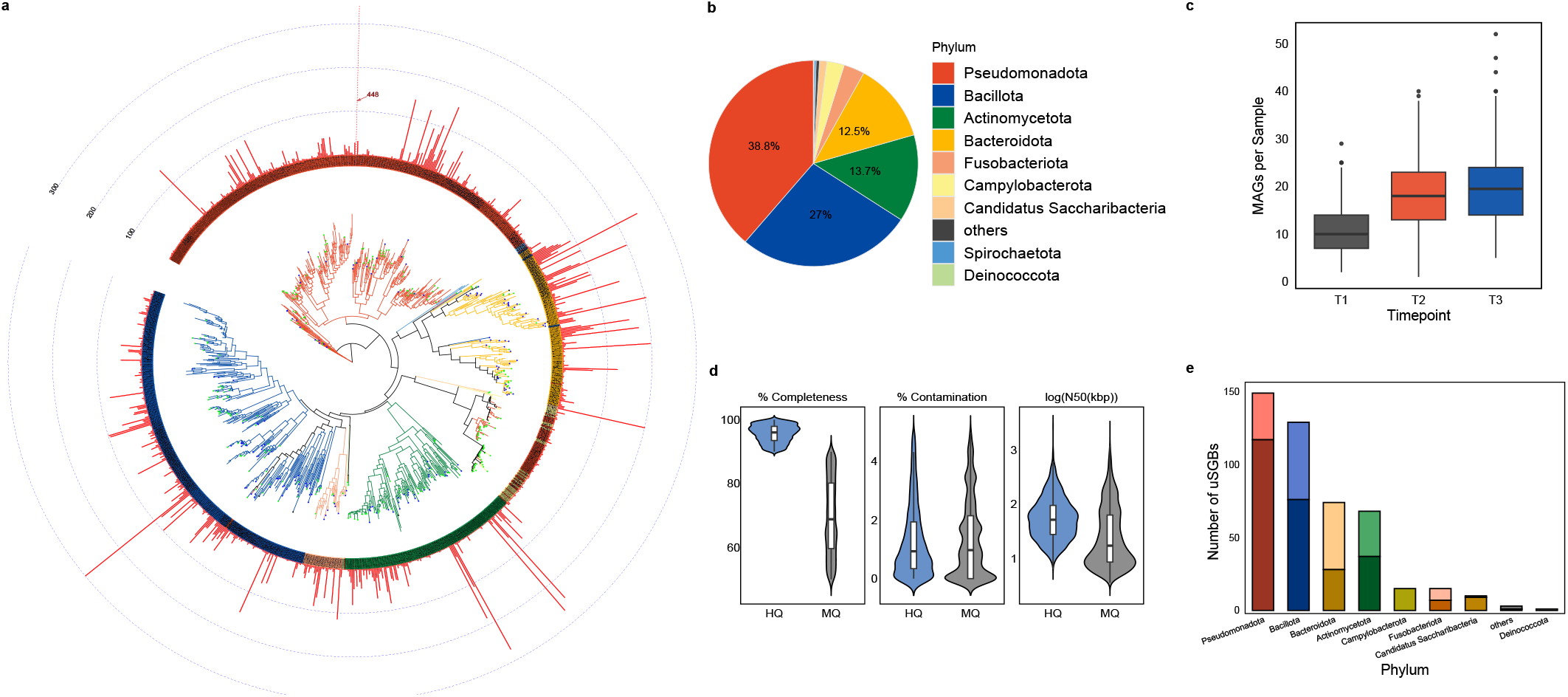
Phylogeny, diversity, and quality of reconstructed MAGs. **a**, Maximum-likelihood phylogenetic tree constructed from representative MAGs using PhyloPhlAn and visualised in iTOL. Tree branches and outer circles are coloured by phylum as shown in **(b)**. Unknown MAGs assigned to genus-level genome bins (GGBs) and family-level genome bins (FGBs) are marked with green and blue stars, respectively. Red bars at the outermost layer represent the number of qualified MAGs belonging to the same bacterial species (i.e. assigned to the same secondary cluster in dRep) that were recovered across all samples. Notably, we identified 26 MAGs belonging to the order *Campylobacterales* within the phylum Campylobacterota (highlighted in yellow), which were highly intermixed with members of the phylum Pseudomonadota (red) in the phylogenetic tree. This pattern reflects the historical classification of *Campylobacterales* under Pseudomonadota in earlier taxonomic systems. The observed phylogenetic proximity likely stems from their high sequence similarity, underscoring the evolutionary continuity between these groups despite recent taxonomic revisions. **b**, Distribution of all qualified MAGs by phylum. **c**, Distribution of qualified MAGs per sample for each timepoint. **d**, Quality statistics of all qualified MAGs stratified into high-quality (HQ) and medium-quality (MQ) categories. **e**, Stacked bar plot showing the distribution of unclassified species-level genome bins (uSGBs) by phylum. For each phylum, the upper bar represents FGBs and the lower bar represents GGBs.

#### Replication rates reveal complex population dynamics in key oral microbial groups

To assess the population dynamics of individual species along oral microbiome development, we estimated microbial population replication rates^31^ from MAGs and compared them with relative abundance. In total, 1,045 MAGs representing 54 species from six key genera, *Corynebacterium, Fusobacterium, Neisseria, Haemophilus, Streptococcus*, and *Actinomyces*, plus 17 MAGs from two species of the phylum Saccharibacteria (also known as TM7) were analysed. All taxa showed replication indices >1, indicating active growth (Fig. 8a-b, Supplementary Table 8).

**Fig. 8:**
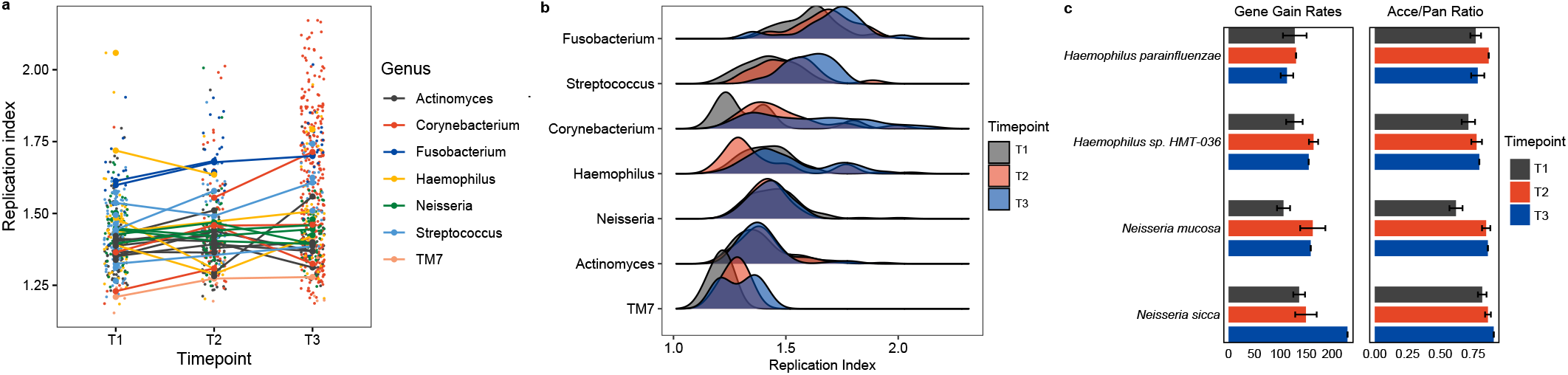
Replication and pangenome dynamics of oral bacteria across childhood. **a**, Replication indices of bacterial species from six major genera and the phylum TM7, estimated using iRep across three timepoints (T1, T2, T3). Lines indicate species-level trajectories by connecting the mean replication index at each timepoint. Points represent replication indices of individual MAGs at each timepoint. **b**, Density plots of replication indices for individual MAGs, with x-axis showing replication index, and y-axis showing bacterial groups. Groups are ordered by mean replication index to highlight group-level differences in replication dynamics. **c**, Pangenome dynamics across development for four MAGs abundant oral species (*Haemophilus parainfluenzae, Haemophilus* sp. HMT-036, *Neisseria mucosa*, and *Neisseria sicca*). Left panels: gene-gain rates, estimated as the slope of the gene accumulation curve from a fitted linear model (genes gained per additional genome) at T1-T3; right panels: accessory-to-pangenome gene ratios at each timepoint based on pangenome analysis in anvi’o. Bars indicate means and error bars show standard deviations from 100 subsampled replicates.

Replication rates varied substantially across species, ranging from 1.15 to 2.17 (Fig. 8a). Despite this heterogeneity, genus-level trends were evident for several taxa, revealing coherent population-level strategies during oral microbiome development. *Fusobacterium* (primarily *F. nucleatum* and *F. pseudoperiodonticum*) consistently exhibited high and increasing replication rates across development (Extended Data Fig. 4), despite maintain relatively low, though rising relative abundance (Pearson r = 0.95). Mean replication rates for Fusobacterium were significantly higher than those of other genera at each timepoint (Welch’s t-test; ANOVA; all p < 0.001). As a well-characterised bridge taxon linking early and late colonisers, this pattern suggests that *Fusobacterium* persistence within the oral microbiome is maintained through sustained growth coupled with high cellular turnover rather than abundance dominance^44,45^. Active replication likely facilitates frequent cross-species interactions and promotes biofilm development across heterogeneous oral niches^32^, consistent with its recognised role in community assembly and metabolic adaptability^33^. Together, these features indicate strong ecological fitness and resilience of *Fusobacterium* within the developing oral environment, while highlighting that abundance-based metrics alone underestimates its ecological impact. Among them, *F. nucleatum* is particularly important, given its established links to oral oncogenesis and its contribution to the pathogenesis of periodontitis^34,35^. It’s population dynamics in early life warrant further attention.

In contrast, *Actinomyces* and *Neisseria* maintained relatively low and stable replication rates while progressively increasing in abundance. This pattern suggests that their ecological success is potentially driven by efficient adhesion to diverse oral niches with reduced cellular turnover and high persistence, consistent with their roles as stable, long-term colonisers during oral microbiome succession. Substantial interspecies variation was observed within *Haemophilus, Streptococcus*, and *Corynebacterium*. All three genera displayed overall increase in replication rates across development. *Haemophilus* and *Corynebacterium* showed concurrent increases in relative abundance, supporting their roles as commensal taxa contributing to biofilm maturation and stability^36^. By contrast, *Streptococcus* exhibited rising replication rates despite declining relative abundance, indicating strong competitive capacity under increasing community complexity as a pioneer coloniser.

#### Pangenome dynamics and functional enrichment of oral bacteria across childhood

We investigated pangenome dynamics of four oral species with abundant MAGs across three developmental stages: *Haemophilus parainfluenzae, Haemophilus* sp. HMT036, *Neisseria mucosa*, and *Neisseria sicca*. To account for uneven sampling, we randomly subsampled MAGs to the minimum group size for each species, then calculated two metrics per timepoint early-stage gene-gain rates and the accessory-to-pangenome ratio (Fig. 8c).

Within‐species comparisons revealed significant effects of timepoints on gene cluster presence profiles (PERMANOVA, all p < 0.05). All four species maintained open pangenomes, indicating ongoing gene acquisition. Gene‐gain rates for *H. parainfluenzae, Haemophilus sp*. HMT036, and *N. mucosa* peaked at T2, while *N. sicca* showed a steady increase to T3. Accessory-to-pangenome ratios rose continuously from T1 to T3 in *Haemophilus* sp. HMT036, *N. mucosa*, and *N. sicca*, while *H. parainfluenzae* again peaked at T2. Functional enrichment of pangenomes further revealed temporal shifts in metabolic capacity. *H. parainfluenzae* exhibited progressive enrichment of histidine and NAD biosynthesis, while fatty-acid biosynthesis and urea-cycle pathways were enriched at T1-T2 but declined by T3. Both *N. sicca* and *N. mucosa* exhibited shared T3‐enriched COG categories, including defence mechanisms, signal transduction, and extracellular structures. Early timepoints in these species were enriched in secondary metabolism, transport and catabolism, cell cycle control, and lipid metabolism. *Haemophilus sp*. HMT036 showed no significant enrichment.

#### Phylogenetic diversity of Saccharibacteria

Species from the phylum Saccharibacteria (TM7) are prevalent in the oral cavity but remain challenging to study due to their symbiotic lifestyle and difficulty in cultivation. Leveraging our large number of samples, we obtained high-quality draft genomes belonging to TM7 and conducted in-depth genomic analyses, including replication rate estimation and phylogenetic reconstruction. TM7 group exhibited the lowest replication rates among all investigated oral bacterial groups, reflecting their symbiotic and parasitic lifestyle that involves reduced metabolic activity^37–39^ (Fig. 8a-b). The gradual increase in replication across childhood suggests progressive adaptation to the oral niche and a potential role in shaping community interactions^40^.

Phylogenomic reconstruction of 613 TM7 MAGs, together with 23 reference and 30 outgroup genomes, revealed four major subgroups (G1, G3, G5, and G6)^40,41^ (Fig. 9). G1 was the largest comprising four monophyletic clades (472 MAGs). The subclade represented by TM7c and TM7x encompassed most oral MAGs, displayed low genetic divergence and was basal within G1, suggesting a conserved, host-adapted lineage. In contrast, the other G1 subclades were represented by environmental references^42,43^ (*Candidatus Microsaccharimonas sossegonensis, Candidatus Saccharimonadaceae aalborgensis, Candidatus Mycosynbacter amalyticus*) and included only a few oral MAGs, reflecting their host and niche specific genomic diversity. Group G6 also exhibited a high degree of internal phylogenetic clustering, with most MAGs related to *Ca. Nanogingivalis bacterium* and *Ca. Saccharimonas* sp., supported by high bootstrap values. Strain *Candidatus Chaera renei* served as the outgroup for G1 and G6, together forming a clade distinct from G3 and G5. Groups G3 and G5 contained far fewer oral MAGs. No MAGs clustered closely with the G5 reference *Ca. Nanoperiomorbus periodonticus* while most G3 MAGs grouped with the reference strain *Ca. Nanosyncoccus nanoralicus*.

**Fig. 9:**
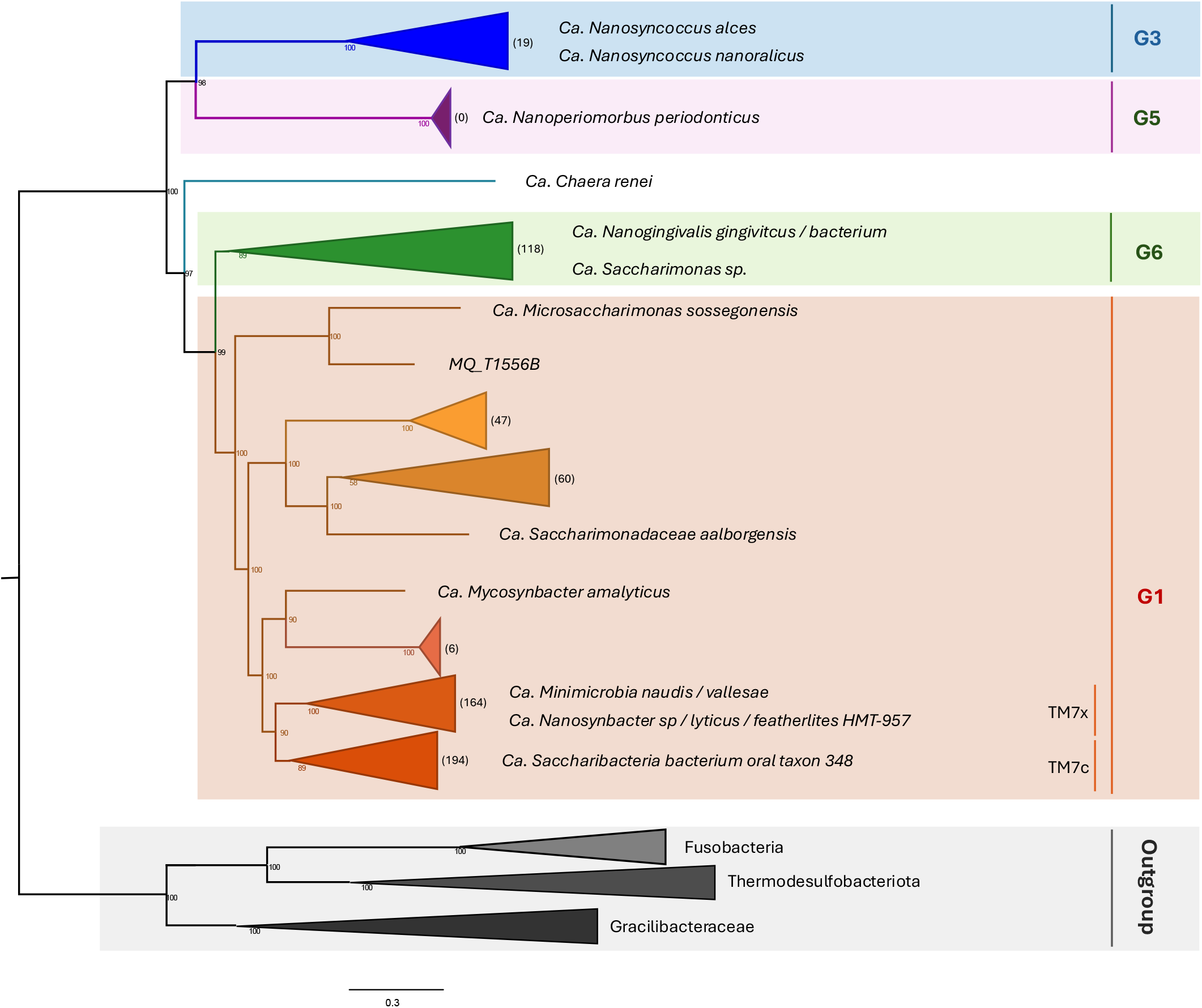
Phylogenomic reconstruction of TM7 MAGs and reference sequences. Maximum-likelihood tree inferred with IQ-TREE from qualified TM7 MAGs reconstructed in this study, together with public references and outgroup genomes. Four major subgroups (G1, G3, G5, G6) were identified and are shown as collapsed and color-coded clades in FigTree. Monophyletic subclades within G1 were also collapsed with TM7c and TM7x subclades forming a basal lineage comprising most oral MAGs. Numbers in brackets beside each subclade indicate the number of recovered MAGs. Bootstrap support values (from 2,000 ultrafast replicates) are shown on tree nodes.

## Discussion

Using a large longitudinal twin cohort, we characterised the developmental trajectory of the oral microbiome across childhood and demonstrated marked age-related changes in both ecological composition and functional capacity. Together with the shifting relative contributions of genetic and environmental influences, these findings provided an integrated picture of oral microbiome development as a dynamic, stage-dependent process. Environmental factors explained most for the variation of oral microbiome during early childhood, consistent with the high variable nature of oral microbiome along an immature immune system. This highlights the sensitivity of early oral colonisation to diet, hygiene, and household exposures, which collectively shape pioneer colonisers and key metabolic pathways. Importantly, the relatively weak influence of host genetics at this stage should be interpreted in comparison to strong environmental effects rather than as an absence of genetic contribution. Although host genetics showed limited association with overall community diversity in early life, substantial heterogeneity in heritability estimates was observed at the level of individual species, underscoring species-specific responses to host and environmental pressures.

By late childhood (T3), oral microbiomes converged toward more similar community profiles while exhibiting increased metabolic versatility and functional complexity. At this stage, heritability emerged as a dominant influence, particularly for pathways related to microbial respiration, nucleic acid synthesis, and energy metabolism. These patterns suggest that maturation of the host oral environment increasingly imposes deterministic constraints on microbial persistence and function. Host genetic effects may act through mechanisms such as immune regulation, salivary composition, and epithelial surface properties; however, microbial–microbial interactions within structured oral biofilms, including metabolic cooperation and signalling, are also likely to play a critical role in shaping these outcomes.

Strain-level variation captures ecologically meaningful diversity that is obscured at the species level, providing deep insight into microbial adaptation, persistence, and host-specific selection. While species identity explained much of the variance in genetic diversity, temporal shifts were evident at the lineage level, indicating species-specific evolutionary trajectories rather than a uniform developmental pattern. In contrast to the human gut, where most species are typically dominated by a single strain with very low polymorphism^44^ (<0.1%) across individuals from diverse backgrounds, oral species in our longitudinal cohort exhibited markedly higher polymorphic rates (0.16-6.87%, Fig. 5b), even when stratified by age. This suggests persistence of multiple genetically distinct lineages and highlights the oral cavity as a dynamic ecological niche shaped by rapid immune surveillance, frequent disturbance from continual exposure to diet, hygiene, and environmental transmission, where evolutionary processes may act more rapidly than in the gut. Many *Streptococcus* species exhibited high genetic diversity and polymorphism during early colonization, reflecting rapid adaptation to host and environmental pressures at different stages. Notably, the cariogenic *S. mutans* showed elevated diversity at T3 and consistently high polymorphism across all timepoints, indicating strong and persistent selection pressures, likely driven by dietary sugars, acid stress, and host immunity^45,46^. This evolutionary flexibility likely contributes to its persistence and pathogenic role in childhood caries.

We refined the phylogeny of TM7 using 666 MAGs and reference genomes spanning major subgroups. Most oral TM7 species in children clustered within G1, reflecting host and niche specific genomic diversity and is consistent with previous studies^40,47^. While earlier studies have reported the prevalence of 4 subgroups (G1, G3, G5, and G6) in humans, we did not recover G5 MAGs. Notably, G5 references were derived from adult patients with severe periodontitis^40^. The phylogenetic placement of TM7 lineages within the oral cavity has remained controversial^40,48^, and our analysis further highlights this complexity by showing a close relationship between G1 and G6 (118 MAGs). These findings suggest that the diversity of TM7 lineages may differ between children and adults, reflecting responses to variation in immune maturation, host environment, or disease status. While we included medium-quality MAGs to expand genomic coverage, the limited number of reference genomes in public databases continues to constrain the classification of additional TM7 lineages. Nevertheless, MAG-based phylogenetic reconstruction provides a powerful approach to resolve TM7 diversity beyond the limits of classical taxonomy and substantially expands the genomic resources available for this unculturable lineage. This study facilitates further exploration of strain-level diversity and the ecological roles of TM7 lineages, including functional differentiation across conditions, symbiotic interactions with host bacteria such as *Actinomyces*, and gene-level diversity involved in particular functions such as for adhesin and stress responses.

In summary, our study demonstrates distinct evolutionary trajectories of the oral microbiome throughout childhood, offering detailed insights into the dynamic host and environmental influences that shape microbial communities during development. Future analyses incorporating additional timepoints and genomic features will provide finer resolution of these dynamics and uncover further aspects of microbial adaptation^49^. Together, these efforts will advance our insights into the oral microbiome’s role in health and disease.

## Methods

### Study design and sample collection

This study was conducted with approval from the Adelaide University Human Research Ethics Committee (H-2013-097 and H-78-2003). Parents provided written informed consent for the collection and use of samples and associated data for research purposes.

The initial dataset comprised 189 families and 380 twin individuals, all Australian children (Supplementary Table 1). A total of 964 oral biofilm samples were collected across three developmental timepoints as described in our previous study^22^, including T1: Edentulous (no teeth), average age 6.8 ± 2.7 months (271 samples, collected from gum); T2: Primary/deciduous (baby teeth only), average age 1.6 ± 0.4 years (313 samples, collected from gum and teeth plaque) and T3: Mixed dentition, average age 8.8 ± 1.2 years (380 samples, collected from supragingival plaque biofilm). Host contaminations were removed by mapping reads against the hg38 human reference genome (GRCh38). The oral biofilm samples exhibited low levels of host (human) DNA contamination, with an average of 14% across all samples. Quality control analysis resulted in the removal of 44 sequencing libraries. Of these, 35 were excluded due to high levels of host DNA contamination (>65% of reads), seven due to anomalous read counts (<1 × 10^7^ or >1 × 10^8^), and two due to active antibiotic use at the time of sampling (Supplementary Table 1).

### Metagenomic sequencing and data pre-processing

DNA was extracted from samples using the DNeasy^®^ PowerSoil^®^ HTP 96 Kit (QIAGEN). All Samples were co-extracted with blanks to monitor for contamination. Metagenomic library preparation followed our previous study^22^ with the Nextera DNA Flex protocol, and sequencing was performed on the Illumina NovaSeq 6000 platform. On average, 62.8 million paired-end reads (134-150 bp) were generated per sample (Supplementary Table 2).

Data pre-processing followed the procedures documented at https://github.com/fancyge/OralMicrobiome/. For each sample, raw reads were quality-checked using FastQC and host contaminants were removed by mapping against the human reference genome (hg38/GRCh38) with BBMap. Samples with >65% host contamination were excluded. Targeted reads that not mapping to the human genome were retained for downstream analyses. Average host contamination was 14% across all samples, with 64% of samples showing less than 10% contamination.

### Reads-based analysis

#### Taxonomic and functional profiling of sequence reads

For taxonomic classification and quantification of the microbial community in each sample, MetaPhlAn3^50^ was employed to map the filtered reads against the maker gene database CHOCOPhlAn_201901. The results were relative abundance of each taxon estimated from the number of reads assigned to the marker genes. A total of 734 bacterial species were obtained across all samples. Function profiling was performed using HUMAnN3^50^ for each sample. This involves mapping the sequencing reads to the reference database ChocoPhlAn to identify and quantify microbial gene families at strain-level resolution. Functional annotations were then assigned using UniRef databases (UniRef50/UniRef90), and metabolic pathways were reconstructed with MetaCyc references. Both unstratified and species-stratified functional profiles were obtained, resulting in 747 unstratified pathways and 2,196 stratified pathways by species after applying a 0.01% abundance filter.

#### Alpha and beta diversity

All analyses were performed in R unless otherwise specified.

We first examined whether there were batch effects raised due to variability in sample preparation or sequencing among different batches of sequences using Principal Component Analysis (PCA) clustering method with taxonomic and functional profiling results using function *prcomp*. Input data was first filtered by excluding 0.01% species, offset by 1 and then normalised using Total Sum Scaling and Centered Log-Ratio (CLR) transformation. Batch effect was found within functional data. A supervised method Partial Least Squares Discriminant Analysis (PLS-DA) was applied to correct batch effects for function data. The corrected table which was already transformed was then used to perform PCoA for functional profiling patterns across timepoints (Fig. 1f).

Alpha diversity was quantified using the Shannon diversity index, calculated with the *diversity* function from package vegan on both taxonomic and functional abundance profiles. Linear mixed-effects model (*lmer*) was applied to test timepoint effect on alpha diversity with fixed effects for timepoint and sex, and a random intercept for family. Significance of fixed effects was evaluated using type II ANOVA (*anova*), and pairwise comparisons between timepoints were performed using *emmeans*, adjusted for multiple testing using Tukey’s.

Beta diversity was assessed using Bray-Curtis distance for bacterial composition abundances (*vegdist*) and Euclidean distance for transformed and batch corrected functional profiles (*dist*). Weighted unifrac distance metrics were computed using a custom script (calculate_unifrac.R) and the MetaPhlAn species tree (mpa_species_tree.nwk), along with species-level abundance profiles. Data were log10-transformed for normalization. Non-metric multidimensional scaling (nMDS) was performed with metaMDS, and ordinations were visualised with ggplot2. Differences by timepoints were tested with PERMANOVA (*adonis2*), adjusting for sex and family effects and accounting for repeated measures within families.

#### Differential abundance analysis and classification models

Differential abundance analysis was performed using MaAsLin2^51^, with timepoint included as a fixed effect and T1, T2, or T3 set as the reference category to enable pairwise comparisons between groups. Species abundance profiles were derived from MetaPhlAn3, and unstratified functional pathway profiles from HUMAnN3 with batch effects corrected. Features were retained for analysis if they had a minimum relative abundance of 0.01% and were present in at least 20% of samples. The analysis used a linear model (LM) framework, applying CLR normalisation with log transformation for taxonomic composition data, while corrected and transformed functional data were analysed without additional normalization.

Classification models were performed using Random Forest algorithms to identify microbial species and functional signatures associated with timepoints using *rf*. The dataset was randomly partitioned into training (75%) and testing (25%) subsets using function *initial_split*, with stratification by timepoint to ensure balanced representation across groups. Models were trained on the training set and evaluated on the test set. Model performance and error rates were estimated across 100 Random Forest runs, reporting the mean and standard deviation. Variable importance was assessed using both MeanDecreaseGini and MeanDecreaseAccuracy to capture complementary aspects of feature contribution. Six datasets were analysed: 1) Relative abundance data of 734 bacterial species generated from MetaPhlAn^50^; 2) A subset of 51 differentially abundant species identified by MaAsLin2^51^; 3) Relative abundance data of 2,196 species-stratified functional pathways from HUMAnN3^50^ (filtered to exclude pathways with < 0.01% abundance, batch effect corrected); 4) A subset of 1,374 differentially abundant species-stratified functional pathways identified by MaAsLin2; 5) Relative abundance data of 425 unstratified pathways, and 6) A set of 229 differentially abundant unstratified pathways identified by MaAsLin2.

#### Univariant ACE structural equation modelling

We fitted univariate ACE structural equation models to decompose phenotypic variance into additive genetic (A), shared environmental (C), and unique environmental (E) components. Models were parameterised using a one-factor Cholesky decomposition implemented in R package *umx*^23,24^. ACE modelling was performed using Shannon diversity as a phenotype representing overall microbiome alpha diversity. For individual species and functional features, stringent filtering criteria were applied first to minimize noise from rare or low-information features prior to modelling. Specifically, for microbial species, only those detected in at least 20% of samples, with mean relative abundance ≥0.001, and variance ≥0.001 after log1p transformation were retained. For functional pathways, we required prevalence ≥20%, mean relative abundance ≥0.001, and removed the bottom quartile of pathways ranked by standard deviation across individuals. To further ensure model stability, features with fewer than 10 complete twin pairs per zygosity group were excluded. After filtering, 147 of 734 species and 181 of 425 functions remained for ACE analysis. For species, MetaPhlAn-derived relative abundances were log1p-transformed prior to ACE modelling. For functional pathways, we used normalised and batch-corrected abundance profiles. To account for covariate effects, we fitted linear models with sex as a covariate, extracted residuals, and standardised them (z-scored) to mean 0 and unit variance. These standardised residuals were used as phenotypic inputs for ACE models. Model fit was compared with nested AE, CE, and E models using likelihood-ratio tests. Confidence intervals of each component were obtained using likelihood-based profile CIs. Intraclass correlation coefficients (ICCs) and 95% confidence intervals were estimated using function *ICC* in R package *psych*.

#### Strain-level analysis

StrainPhlAn3 was employed for strain-level profiling of microbial communities which relies on a set of informative species-specific marker genes to reconstruct the dominant strain in each sample^44^. Forty-four species were selected based on abundance (Supplementary Table 9), differential abundance signals from MaAsLin2, and important abundant genera in the oral biofilm (*Streptococcus, Neisseria, Veillonella*). For each sample, the target reads were first mapped against the MetaPhlAn markers by Bowtie2 to generate the consensus sequence, which represented the most abundant strain per species. The consensus sequences for public reference genome strains were also generated by aligning marker genes to reference genomes. These consensus sequences from both references and samples were then aligned using MUSCLE, and phylogenetic trees were constructed using RAxML (-m GTRCAT and -p 1234).

Multiple‐sequence alignments containing fewer than five sequences at any timepoint were excluded to avoid unreliable distance estimates. Consequently, *Veillonella sp*. T11011_6 and *Streptococcus pseudopneumoniae* were omitted due to insufficient coverage. For each retained species, pairwise genetic distances (1.9 million comparisons across 44 species) were calculated using multiple sequence alignments from StrainPhlAn with function *dist*.*alignment*, which measured as the square root of the proportion of mismatched positions.

Polymorphic site was deemed polymorphic if the dominant allele frequency was below 80% in its marker gene alignment. Polymorphic rates were reported by scanning each marker gene to capture genetic dynamics within microbial populations and was defined as the median percentage of such sites across all marker genes for each species. Distance matrices were also generated using *dismat* from EMBOSS with the Kimura nucleotide substitution model, and principal coordinates analysis (PCoA) was performed with *cmdscale*. To test the effects of species, timepoint, and their interaction on strain-level diversity, we applied two-way ANOVA. We further validated species-specific versus global effects with a linear mixed model including species as a random effect. Finally, temporal trends were assessed separately for each species using one-way ANOVA, with *p*-values adjusted for multiple testing using the Benjamini-Hochberg false discovery rate (FDR).

To test whether within-species genetic variation differed across timepoints for *S. mitis*, we applied both PERMANOVA (*adonis2*, 1,000 permutations) and ANOSIM (*anosim*, 1,000 permutations) as an additional rank-based test of timepoint separation. The results of statistic R and p-value were used for interpretation where a higher R meaning more dissimilarity between groups. To account for potential bias due to differences in within-group variability, homogeneity of multivariate dispersion was tested using *betadisper* with both ANOVA and permutation-based tests (*permutest*).

#### Gene-content profiling

PanPhlAn was applied to perform strain-level metagenomic profiling by identifying the species-specific gene composition within individual metagenomic samples for *S. mitis*. Metagenomic reads were mapped against the pan-genome of *S. mitis* to infer the presence of core and accessory genes in each sample.

To control for unequal sample sizes across timepoints in the result (T1 = 152, T2 = 81, T3 = 63), all richness and overlap analyses employed balanced subsampling. Each timepoint was repeatedly subsampled to 60 samples across 1,000 iterations. Gene family and marker gene richness were recalculated to obtain mean estimates and 95% confidence intervals. Gene families detected in fewer than five samples were excluded to avoid convergence issues. For each remaining gene family, we fit a logistic regression model with presence/absence as the outcome and sampling timepoint as the predictor to assess whether timepoint significantly explained variation in gene family presence. Resulting p-values were adjusted for multiple testing using FDR correction, and gene families with an adjusted p-value < 0.05 were considered significantly variable across timepoints.

Upset plot of gene-family distributions across timepoints were plotted using gene-content profiling with function *upset*. Set size was derived from the balanced subsampling procedure, and intersection sizes represent the median values across 1000 subsampled iterations.

### Metagenome-assembled genomes (MAGs) construction and analysis

#### Metagenome assembly and MAGs reconstruction

For each sample, target reads were assembled into contigs using MEGAHIT with default parameters. Paired-end reads were then aligned back to the assemblies using BWA-MEM, and coverage profiles were generated with Samtools and Sambamba. Contigs were subsequently binned into metagenome-assembled genomes (MAGs) using MetaBAT2^52^ with default settings, incorporating coverage information. The quality of each draft genome was examined using tool CheckM in terms of completeness and contamination rates. MAGs were categorized into high-quality (completeness ≥ 90%; contamination < 5%) and medium-quality (completeness ≥ 50% and completeness < 90%; contamination < 5%) genomes based on the MIMAG standard^28^ with minor modification.

#### Dereplication of MAGs and Taxonomy assignment of SGBs

A total of 15,393 MAGs meeting medium and high-quality criteria underwent further analysis. Dereplication of these qualified MAGs was performed using dRep^29^, resulting in 965 species-level genome bins (SGBs) (settings: -comp 75, -pa 0.9, -sa 0.95). Taxonomic assignments for the 965 species-level genome bins (SGBs) were conducted using FastANI^53^ and Mash^54^, leveraging their complementary approaches to genome similarity assessment. Specifically, FastANI was first applied to compare all SGBs against complete reference genomes from both NCBI RefSeq and GenBank (as of Mar 2025), providing high-resolution assessments of sequence identity. FastANI only reports matches with ANI (Average Nucleotide Identity) above 70%, which resulted in successful matches for 889 MAGs. To further validate and supplement these results, Mash was employed to compare MinHash sketches of all the MAGs against two reference databases: one constructed from the same set of complete reference genomes as above, and another based on the precomputed Mash sketch database from RefSeq release 88. Notably, RefSeq release 88 includes both complete and incomplete genomes, allowing Mash to detect potential novel sequences that may have been missed using only complete references. Due to their methodological differences, FastANI, which requires high-quality, near-complete genomes, provides precise ANI calculations but can be sensitive to recombination and horizontal gene transfer. In contrast, Mash, which relies on k-mer similarity, offers greater flexibility in detecting distant relationships, although with less specificity.

Taxonomic assignments were determined based on the accession number of the closest reference genome identified using either FastANI or Mash. MAGs were assigned to known species-level genome bins (kSGBs) when the maximum ANI from FastANI was ≥ 95%, identifying 404 kSGBs. Additionally, MAGs with ANI values between 90% and 95% and a minimum Mash distance ≤ 0.05 were assigned to putative kSGBs; MAGs with completeness > 90% and a minimum Mash distance ≤ 0.05 were also considered for putative kSGB assignment, resulting in a total of 92 additional kSGBs (Supplementary Table 7). The remaining 469 unclassified species-level genome bins (uSGBs) were designated as novel species, as they showed no species-level matches in available reference databases and exhibited ANI values between 74.5% and 94.9%. To further resolve their taxonomy, a hierarchical classification strategy was applied. uSGBs were assigned to genus-level genome bins (GGBs) when they showed a maximum ANI of 85-95% or a minimum Mash distance between 0.05 and 0.34^55^. Similarly, those with a maximum ANI < 85% or Mash distance > 0.34 were classified into family-level genome bins (FGBs). As a result, 290 SGBs were assigned to GGBs, 174 to FGBs and 5 remained unassigned. Given the resolution limits of these methods, taxonomic assignments for GGBs and FGBs are considered reliable only at the genus and family levels, respectively.

#### Replication rates analysis

Replication rates were estimated using tool iRep^31^ with default settings (min cov = 5, min wins = 0.98, min r^2^ = 0.9, max fragments/Mbp = 175, GC correction min r^2^ = 0.0). To select MAGs of a particular species, we first identified the highest quality representative MAG of each species from dereplicated genomes from tool dRep. All other qualified MAGs belonging to the same secondary cluster as the representative MAGs were then collected. These MAGs were further filtered to retain only those with ≥ 75% completeness, ≤ 175 fragments/Mbp sequence and ≤ 2% contamination. For TM7, selection criteria were slightly relaxed due to the limited number of publicly available reference and the reduced genome sizes with fewer marker genes, which could otherwise lead to underestimation of sequence quality. Specifically, we maintained the criteria but additionally included 4 MAGs from T1 samples that had slightly lower completeness (≥ 65%) but 0% contamination. To evaluate whether replication rates of Fusobacterium were significantly higher than the other taxa, we first collapsed genera into two groups: *Fusobacterium* and non-*Fusobacterium*. Replication indices were compared between the two groups using Welch’s two-sample *t*-test to account for unequal variances for each timepoint and further assessed using a one-way ANOVA. To test whether replication rates varied across all selected genera, we performed a separate one-way ANOVA including all genus-level groups. Correlation of dynamics of replication rates and relative abundances of selected groups were assessed using Pearson method with function *cor*. To compare distributions across groups and timepoints, we generated density ridge plots using package *ggridges*.

#### Pangenome construction and functional enrichment of MAGs

Pangenomes were built with anvi’o^56^ by selecting MAGs with ≥ 70% completeness and ≤ 5% contamination rate, retaining species with ≥ 8 MAGs per timepoint and ≤ 200 total MAGs. To compare pangenome openness across development, we subsampled each species to its minimum per-timepoint MAG count and computed the early gene-accumulation slope (gene-gain rate) over the first 8 genomes. The accessory‐to‐pangenome ratio was calculated as the number of accessory gene clusters divided by total pangenome gene clusters. All calculation were normalized by subsampling same number of MAGs for each time point and repeated 100 times per species. Functional enrichments were analysed for each species across timepoints using COG20 annotations. Significant enrichments were obtained with adjusted q value ≤ 0.05 and enrichment score ≥ 2. PERMANOVA tests were applied to test the effects of timepoint using presence absence profiles.

#### Phylogeny reconstruction

Phylogeny of 965 representative MAGs were reconstructed using PhyloPhlAn^57^ with a set of 400 universal marker genes with parameter (--min_num_markers 50). The final consensus alignment comprised of 1,783 amino acid variant sites from 936 genomes. Maximum likelihood inference was performed using RAxML under the PROTCATLG model. The resulting tree was visualised in iTOL^58^, with phylum-level taxonomy and abundance information annotated.

For the TM7 clade, we analysed 613 MAGs classified within this phylum (77 known MAGs and 536 unknown MAGs from qualified MAGs), together with 23 reference genomes from public databases that covering 4 well-known major subgroups of TM7 and 30 outgroup genomes (10 each from Thermodesulfobacteriota, Gracilibacteraceae, and Fusobacteria; Supplementary Table 10) to improve phylogenetic resolution. Multiple sequence alignments were generated using PhyloPhlAn with a minimum of 50 markers per genome. Phylogeny was inferred using IQ-TREE under the LG+F+G+C10 model with 2000 ultrafast bootstrap replicates (-bb 2000). The final alignment included an average of 92 marker genes across 651 genomes, comprising 2,060 amino acid variant sites. Tree visualization and clade annotation were performed using FigTree.

## Data availability

The sequencing data generated in this study have been deposited in the European Nucleotide Archive under accession number PRJEB54673 and PRJEB98153.

## Codes availability

Codes for bioinformatics pipelines and statistics are available at https://github.com/fancyge/OralMicrobiome/.

## Authors contribution

F.W. and C.J.A. conceived and designed the study. F.W. developed the methodology, performed the bioinformatics and statistical analyses, data visualisation, and writing. A.J.H. contributed to manuscript critical reviewing and editing. G.V.B. and C.J.A. conducted the laboratory work. M.R.B. and K.M.D. contributed to sample collection and management. T.E.H. assisted with sample curation and study coordination. X.H. contributed to manuscript reviewing. C.J.A contributed to reviewing and administered the project.

## Acknowledgement

We thank all participating twins and their families, as well as the dental staff involved in oral sample collection. We also acknowledge the high-performance computing resources provided by the Australia’s National Computational Infrastructure and the University of Sydney high-performance computing cluster. This work was supported by the National Health and Medical Research Council (APP1062911) to C.J.A and the National Institutes of Health grant (DE029838) to C.J.A.

## References

1. Gomez, A. & Nelson, K. E. The Oral Microbiome of Children: Development, Disease, and Implications Beyond Oral Health. Microb. Ecol. 73, 492–503 (2017).

2. Könönen, E. Development of oral bacterial flora in young children. Ann. Med. 32, 107–112 (2000).

3. Sampaio-Maia, B. & Monteiro-Silva, F. Acquisition and maturation of oral microbiome throughout childhood: An update. Dent. Res. J. (Isfahan). 11, 291–301 (2014).

4. Caufield, P. W. et al. Natural history of Streptococcus sanguinis in the oral cavity of infants: evidence for a discrete window of infectivity. Infect. Immun. 68, 4018–4023 (2000).

5. Tonelli, A., Lumngwena, E. N. & Ntusi, N. A. B. The oral microbiome in the pathophysiology of cardiovascular disease. Nat. Rev. Cardiol. 20, 386–403 (2023).

6. Xu, Q. et al. The oral-gut microbiota axis: a link in cardiometabolic diseases. NPJ Biofilms Microbiomes 11, 11 (2025).

7. Baker, J. L., Mark Welch, J. L., Kauffman, K. M., McLean, J. S. & He, X. The oral microbiome: diversity, biogeography and human health. Nat. Rev. Microbiol. 22, 89–104 (2024).

8. Gomez, A. et al. Host Genetic Control of the Oral Microbiome in Health and Disease. Cell Host Microbe 22, 269–278.e3 (2017).

9. Blekhman, R. et al. Host genetic variation impacts microbiome composition across human body sites. Genome Biol. 16, 191 (2015).

10. Adler, C. J., Cao, K.-A. L., Hughes, T., Kumar, P. & Austin, C. How does the early life environment influence the oral microbiome and determine oral health outcomes in childhood? Bioessays 43, e2000314 (2021).

11. Sitarik, A. R. et al. Fetal and early postnatal lead exposure measured in teeth associates with infant gut microbiota. Environ. Int. 144, 106062 (2020).

12. Adler, C. J. et al. Sequencing ancient calcified dental plaque shows changes in oral microbiota with dietary shifts of the Neolithic and Industrial revolutions. Nat. Genet. 45, 450–455 (2013).

13. Goodrich, J. K. et al. Genetic Determinants of the Gut Microbiome in UK Twins. Cell Host Microbe 19, 731–743 (2016).

14. Demmitt, B. A. et al. Genetic influences on the human oral microbiome. BMC Genomics 18, 659 (2017).

15. Gomez, A. et al. Host Genetic Control of the Oral Microbiome in Health and Disease. Cell Host Microbe 22, 269–278.e3 (2017).

16. Freire, M. et al. Longitudinal Study of Oral Microbiome Variation in Twins. Sci. Rep. 10, 1–10 (2020).

17. Fackelmann, G. et al. Gut microbiome signatures of vegan, vegetarian and omnivore diets and associated health outcomes across 21,561 individuals. Nat. Microbiol. 10, 41–52 (2025).

18. Kaelin, E. A. et al. Longitudinal gut virome analysis identifies specific viral signatures that precede necrotizing enterocolitis onset in preterm infants. Nat. Microbiol. 7, 653–662 (2022).

19. Kageyama, S. & Takeshita, T. Development and establishment of oral microbiota in early life. J. Oral Biosci. 66, 300–303 (2024).

20. Eriksen, C. et al. Early life factors and oral microbial signatures define the risk of caries in a Swedish cohort of preschool children. Sci. Rep. 14, 8463 (2024).

21. Crielaard, W. et al. Exploring the oral microbiota of children at various developmental stages of their dentition in the relation to their oral health. BMC Med. Genomics 4, 22 (2011).

22. Sukumar, S. et al. Development of the oral resistome during the first decade of life. Nat. Commun. 14, 1291 (2023).

23. Rijsdijk, F. V & Sham, P. C. Analytic approaches to twin data using structural equation models. Brief. Bioinform. 3, 119–133 (2002).

24. Bates, T. C., Maes, H. & Neale, M. C. umx: Twin and Path-Based Structural Equation Modeling in R. Twin Res. Hum. Genet. 22, 27–41 (2019).

25. Desai, M. M. & Plotkin, J. B. The polymorphism frequency spectrum of finitely many sites under selection. Genetics 180, 2175–2191 (2008).

26. Karki, R., Pandya, D., Elston, R. C. & Ferlini, C. Defining “mutation” and “polymorphism” in the era of personal genomics. BMC Med. Genomics 8, 37 (2015).

27. Welch, J. J. The Creativity of Natural Selection and the Creativity of Organisms: Their Roles in Traditional Evolutionary Theory and Some Proposed Extensions BT - Evolutionary Biology: Contemporary and Historical Reflections Upon Core Theory. in (eds. Dickins, T. E. & Dickins, B. J. A.) 65–107 (Springer International Publishing, Cham, 2023). doi:10.1007/978-3-031-22028-9_5.

28. Bowers, R. M. et al. Minimum information about a single amplified genome (MISAG) and a metagenome-assembled genome (MIMAG) of bacteria and archaea. Nat. Biotechnol. 35, 725–731 (2017).

29. Olm, M. R., Brown, C. T., Brooks, B. & Banfield, J. F. dRep: a tool for fast and accurate genomic comparisons that enables improved genome recovery from metagenomes through de-replication. ISME J. 11, 2864–2868 (2017).

30. Mirete, S. et al. Metagenome-Assembled Genomes (MAGs): Advances, Challenges, and Ecological Insights. Microorganisms vol. 13 Preprint at 10.3390/microorganisms13050985 (2025).

31. Brown, C. T., Olm, M. R., Thomas, B. C. & Banfield, J. F. Measurement of bacterial replication rates in microbial communities. Nat. Biotechnol. 34, 1256–1263 (2016).

32. Brennan, C. A. & Garrett, W. S. Fusobacterium nucleatum — symbiont, opportunist and oncobacterium. Nat. Rev. Microbiol. 17, 156–166 (2019).

33. Krieger, M. et al. Stratification of Fusobacterium nucleatum by local health status in the oral cavity defines its subspecies disease association. Cell Host Microbe 32, 479–488.e4 (2024).

34. Groeger, S., Zhou, Y., Ruf, S. & Meyle, J. Pathogenic Mechanisms of Fusobacterium nucleatum on Oral Epithelial Cells. Frontiers in Oral Health 3, (2022).

35. Chiscuzzu, F. et al. Current Evidence on the Relation Between Microbiota and Oral Cancer-The Role of Fusobacterium nucleatum-A Narrative Review. Cancers (Basel). 17, (2025).

36. Kreth, J., Helliwell, E., Treerat, P. & Merritt, J. Molecular commensalism: how oral corynebacteria and their extracellular membrane vesicles shape microbiome interactions. Frontiers in oral health 5, 1410786 (2024).

37. He, X. et al. Cultivation of a human-associated TM7 phylotype reveals a reduced genome and epibiotic parasitic lifestyle. Proc. Natl. Acad. Sci. U. S. A. 112, 244–249 (2015).

38. Marcy, Y. et al. Dissecting biological “dark matter” with single-cell genetic analysis of rare and uncultivated TM7 microbes from the human mouth. Proceedings of the National Academy of Sciences 104, 11889–11894 (2007).

39. S., K. R. et al. Small Genomes and Sparse Metabolisms of Sediment-Associated Bacteria from Four Candidate Phyla. mBio 4, 10.1128/mbio.00708-13 (2013).

40. McLean, J. S. et al. Acquisition and Adaptation of Ultra-small Parasitic Reduced Genome Bacteria to Mammalian Hosts. Cell Rep. 32, 107939 (2020).

41. Bor, B., Bedree, J. K., Shi, W., McLean, J. S. & He, X. Saccharibacteria (TM7) in the Human Oral Microbiome. J. Dent. Res. 98, 500–509 (2019).

42. Batinovic, S., Rose, J. J. A., Ratcliffe, J., Seviour, R. J. & Petrovski, S. Cocultivation of an ultrasmall environmental parasitic bacterium with lytic ability against bacteria associated with wastewater foams. Nat. Microbiol. 6, 703–711 (2021).

43. Albertsen, M. et al. Genome sequences of rare, uncultured bacteria obtained by differential coverage binning of multiple metagenomes. Nat. Biotechnol. 31, 533–538 (2013).

44. Truong, D. T., Tett, A., Pasolli, E., Huttenhower, C. & Segata, N. Microbial strain-level population structure & genetic diversity from metagenomes. Genome Res. 27, 626–638 (2017).

45. Bradshaw, D. J. & Lynch, R. J. M. Diet and the microbial aetiology of dental caries: new paradigms. Int. Dent. J. 63 Suppl 2, 64–72 (2013).

46. Oli, M. W., Rhodin, N., McArthur, W. P. & Brady, L. J. Redirecting the humoral immune response against Streptococcus mutans antigen P1 with monoclonal antibodies. Infect. Immun. 72, 6951–6960 (2004).

47. Shaiber, A. et al. Functional and genetic markers of niche partitioning among enigmatic members of the human oral microbiome. Genome Biol. 21, 292 (2020).

48. Murugkar, P. P., Collins, A. J., Chen, T. & Dewhirst, F. E. Isolation and cultivation of candidate phyla radiation Saccharibacteria (TM7) bacteria in coculture with bacterial hosts. J. Oral Microbiol. 12, 1814666 (2020).

49. Wang, F. & Tekle, Y. I. Variation of natural selection in the Amoebozoa reveals heterogeneity across the phylogeny and adaptive evolution in diverse lineages. Front. Ecol. Evol. 10, 1–17 (2022).

50. Beghini, F. et al. Integrating taxonomic, functional, and strain-level profiling of diverse microbial communities with biobakery 3. Elife 10, 1–42 (2021).

51. Mallick, H. et al. Multivariable association discovery in population-scale meta-omics studies. PLoS Comput. Biol. 17, e1009442 (2021).

52. Kang, D. D. et al. MetaBAT 2: an adaptive binning algorithm for robust and efficient genome reconstruction from metagenome assemblies. PeerJ 7, e7359 (2019).

53. Jain, C., Rodriguez-R, L. M., Phillippy, A. M., Konstantinidis, K. T. & Aluru, S. High throughput ANI analysis of 90K prokaryotic genomes reveals clear species boundaries. Nat. Commun. 9, 5114 (2018).

54. Ondov, B. D. et al. Mash: fast genome and metagenome distance estimation using MinHash. Genome Biol. 17, 132 (2016).

55. Zhu, Z., Ren, J., Michail, S. & Sun, F. MicroPro: using metagenomic unmapped reads to provide insights into human microbiota and disease associations. Genome Biol. 20, 154 (2019).

56. Eren, A. M. et al. Anvi’o: An advanced analysis and visualization platformfor ‘omics data. PeerJ 2015, 1–29 (2015).

57. Asnicar, F. et al. Precise phylogenetic analysis of microbial isolates and genomes from metagenomes using PhyloPhlAn 3.0. Nat. Commun. 11, 2500 (2020).

58. Letunic, I. & Bork, P. Interactive Tree Of Life (iTOL) v5: an online tool for phylogenetic tree display and annotation. Nucleic Acids Res. 49, W293–W296 (2021).

