## SupplementaryData for "Ecological and evolutionary dynamics of the oral microbiome across childhood"

### Supplementary data

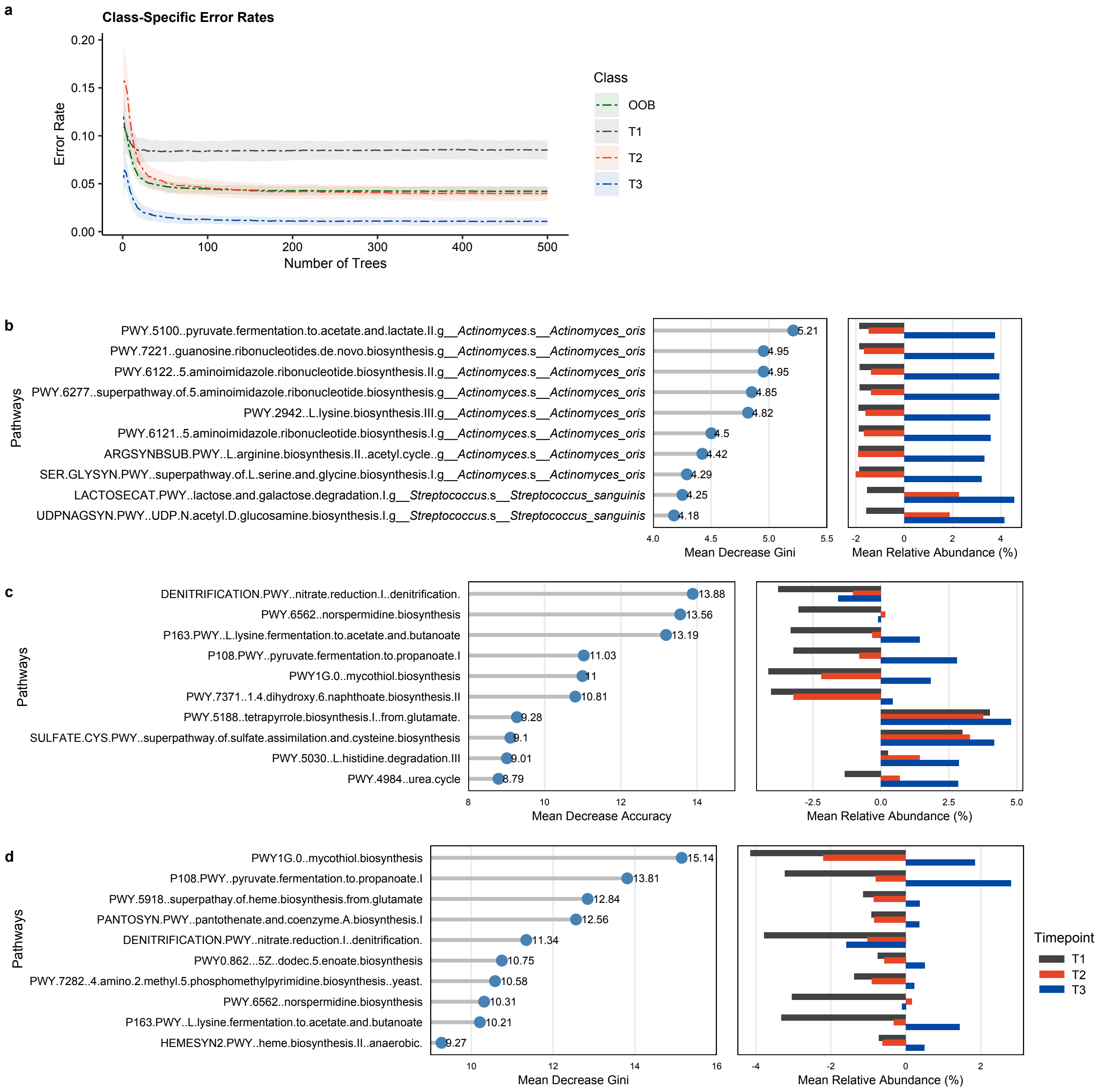


**Extended Data Fig. 1. Key pathway features associated with timepoints identified by random forest models**. **a**, Error rate plot showing the mean and standard deviation across 100 random forest runs, based on abundance profiles of 2196 species-stratified pathways (75% training, 25% testing; split stratified by timepoint; 500 trees per model). OOB: out-of-bag error estimate from internal cross-validation within the training set. **b**, Top ten species-stratified pathways ranked by Mean Decrease Gini based on their abundance profiles. Left panel: Mean Decrease Accuracy for each pathway; right panel: mean relative abundance across samples per timepoint. **c-d**, Top ten unstratified pathways ranked by Mean Decrease Accuracy (**c**) and Mean Decrease Gini (**d**), based on abundance profiles of unstratified pathways

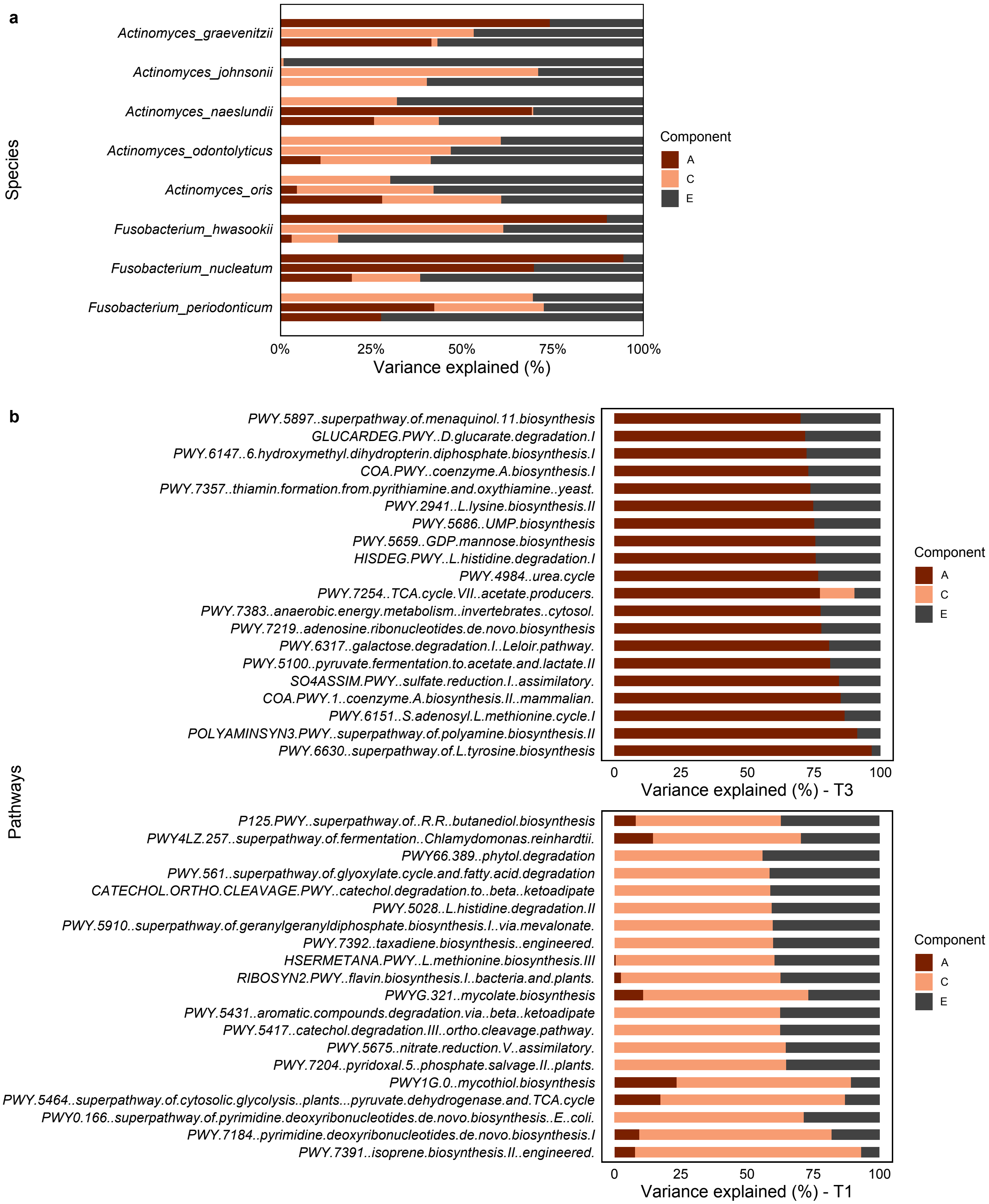


**Extended Data Fig. 2. Partitioning of host genetics and environmental contributions to variations of species and pathways.**

**a,** relative contributions to *Actinomyces* and *Fusobacterium* species from T1 (top) to T3 (bottom). Species are ordered vertically according to the proportion of additive genetic or shared environmental contributions**; b,** Top: Pathways most strongly influenced by host genetics at T3. Bottom: Pathways primarily shaped by unique environmental factors at T1. A = additive genetic, C = shared environmental, E = unique environmental.

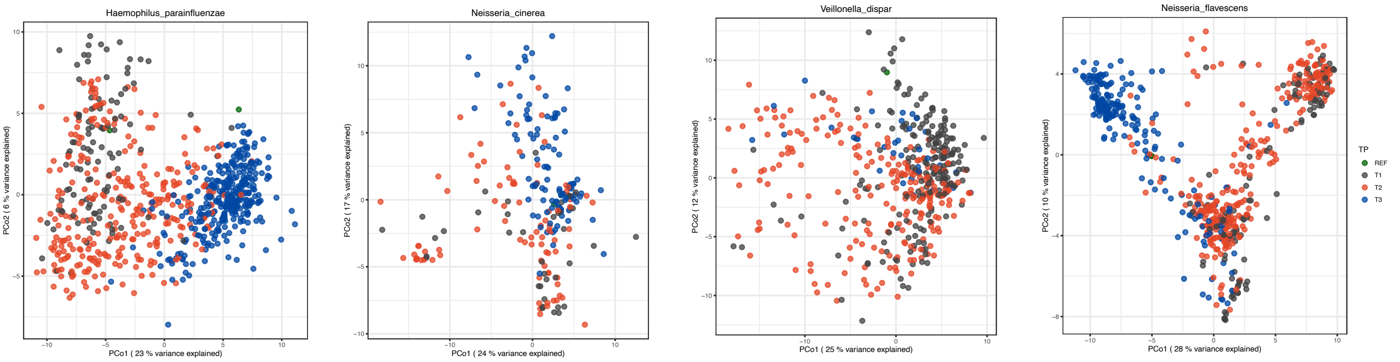


**Extended Data Fig. 3. Genetic diversity of strain-level variation across timepoints.**
Principal Coordinates Analysis (PCoA) based on pairwise genetic distances derived from StrainPhlAn multiple-sequence alignments for four species with the highest variance explained by timepoint, demonstrating clear separation of species populations across developmental timepoints.

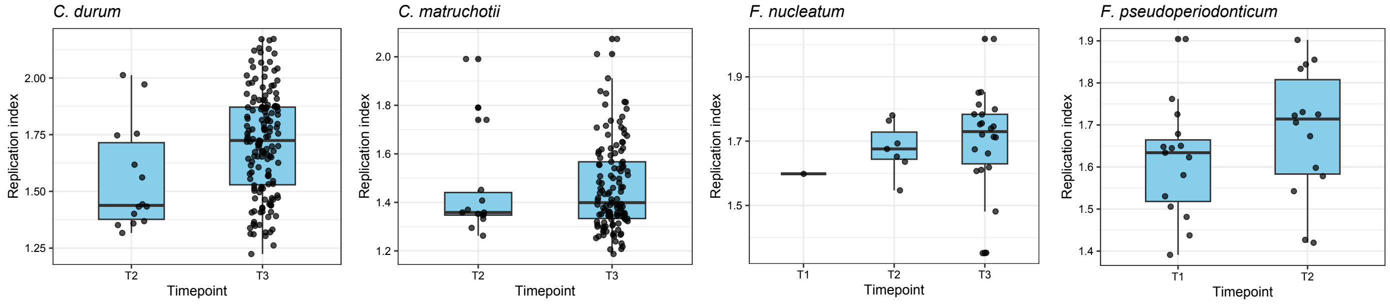


**Extended Data Fig. 4. Replication rates of selected species across timepoints.**
Boxplots display the median and IQR (25th–75th percentiles), with whiskers extending to 1.5× IQR. Points represent replication indices of individual MAGs for each species at each timepoint.

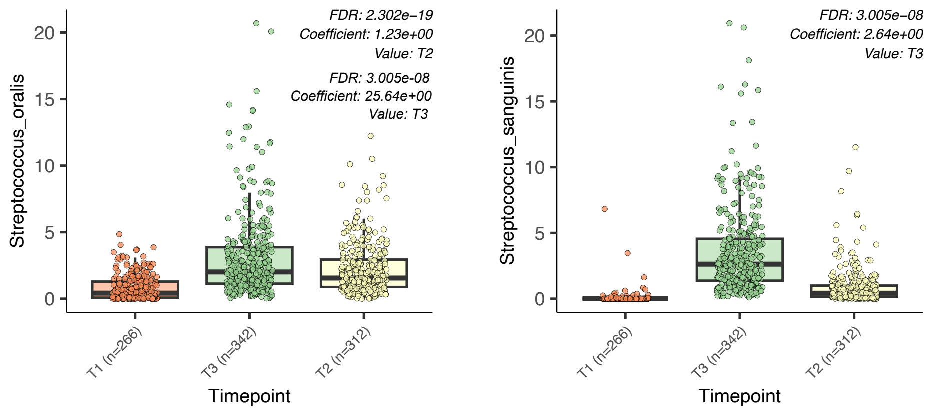


**Extended Data Fig. 5. Differential abundance of *Streptococcus oralis* and *Streptococcus sanguinis* across three timepoints, showing a significant increase at T3.** Results are generated from MaAslin2.

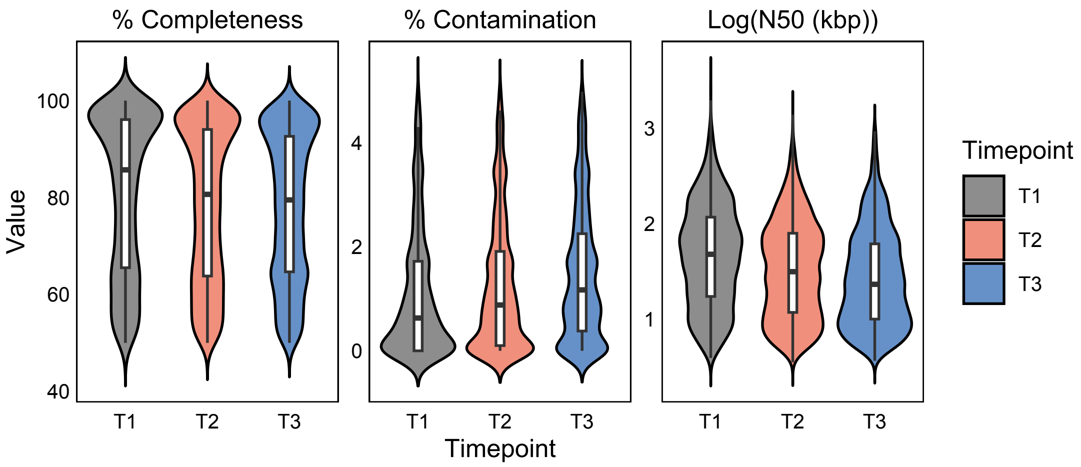


**Extended Data Fig. 6. Quality of MAGs across timepoints.**

### Supplementary Results and Discussions

### Results

#### Differential abundance analysis

Differential abundance analysis identified 51 species as significantly different when comparing across three timepoints (MaAsLin2, Supplementary Table 11). Among them, nine were *Streptococcus* species, most of which displayed significant decrease in relative abundance from T1 to T3. In contrast, *Streptococcus oralis* and *Streptococcus sanguinis* increased significantly as children grew (Extended Data Fig. 5), suggesting their potential protective roles in stabilizing the oral microbiome and counteracting pathogenic Streptococcus species during maturation[1].

A total of 229 pathways with differential abundances were identified, including 179 enriched in T3 (Supplementary Table 12). Notable enriched functions at T3 include tetrapyrrole biosynthesis I from glutamate (PWY-5188), and 5-aminoimidazole ribonucleotide biosynthesis II (PWY-6122, PWY-6277). Note that PWY-6122 and PWY-6277 pathways generate key intermediates for purine nucleotide and thiamine biosynthesis which are essential for energy metabolism[2]. Their abundance has been linked to intestinal diseases and metabolic conditions such as inflammatory bowel disease and hyperuricemia, highlighting their role in host-microbe interactions and inflammatory responses[3,4]. Their enrichment in later childhood oral microbiomes may reflect the maturation of metabolically versatile communities with potential implications for both local oral health and systemic immune modulation in later life.

#### Functional diversity in children oral microbiome

For the overall community, relative abundance analysis showed that the predominant functional capabilities (top 10) remained consistent across T1, T2, and T3, including pathways for nucleotide biosynthesis (adenosine and guanosine), cell-wall construction, and branched-chain amino acid production (L-valine and L-isoleucine). These functions represent core bacterial processes essential for cellular maintenance and survival as expected, and their consistent detection supports the robustness of the dataset.

Functional pathways that were identified as predictive from classification model were linked to *S. peroris*, *S. sanguinis*, *A. oris* (Fig. 3e). These included lactose and galactose degradation I (LACTOSECAT-PWY), purine ribonucleosides degradation (PWY0-1296), L valine biosynthesis (VALSYN-PWY), UDP-N-acetyl-D-glucosamine biosynthesis I (UDPNAGSYN-PWY) and peptidoglycan maturation (PWY0-1586).

### Discussions

#### Microorganisms within oral microbiome

Although the oral microbiome comprises a diverse array of microorganisms, including fungi, archaea, viruses, and protozoa, this study specifically focused on the bacterial component. Non-bacterial taxa typically exist at significantly lower relative abundances compared to bacteria and consequently, they often fall below the detection sensitivity of marker-gene profiling tools like MetaPhlAn3. Furthermore, the DNA-specific extraction protocol employed in this study inherently precludes the recovery of RNA viruses, which constitute a significant portion of the oral virome.

We acknowledge that sampling sites within the oral can have impact in the oral ecosystem. In our longitudinal design, the sampling sites differed across timepoints (gingival samples at T1 versus supragingival plaque at T3, and a combination at T2). Oral niches such as gingival mucosa and tooth surfaces host distinct communities in adults due to differences in adhesion substrates, oxygen exposure, and plaque maturity[5]. However, in early childhood, plaque biofilms are relatively sparse and less structured, and the continuous flow of saliva promotes frequent microbial exchange across niches. Consequently, microbial communities in the oral of young children are more homogenized across different sites than adults and our sampling strategy provides a comprehensive representation of the longitudinal development of the oral microbiome. The use of supragingival samples was particularly informative, as they are more closely associated with caries status and therefore provide relevant insights for understanding caries risk and informing potential management strategies.

**Reads and MAGs in the metagenomic analysis**

We analysed the oral microbiome using both read-based profiling and metagenome-assembled genomes (MAGs) to achieve a comprehensive view of community dynamics. Read-based profiling was used for abundance estimation, as it captures the full community signal by mapping sequencing reads to clade-specific marker genes using MetaPhlAn3. In parallel, taxonomic assignment of representative MAGs was performed using FastANI and Mash, with comparisons against public genomes from the NCBI and RefSeq reference databases.

Overall, the taxonomy recovered from MAGs was broadly consistent with read-based profiles. MetaPhlAn3 identified 734 species from reads, while we recovered 965 dereplicated representative MAGs from all qualified genome bins. This discrepancy reflects methodological differences that read-based approaches capture all detectable taxa present in sequencing reads, whereas MAG-based analyses depend on successful assembly and binning of genomes, which are influenced by community complexity and strain heterogeneity. Note that MetaPhlAn reports relative abundances only for taxa represented by clade-specific marker genes in its reference database and does not include an explicit "unclassified" category; therefore, reads that cannot be assigned to known markers are excluded from taxonomic profiles, leading to underrepresentation of some species. Expected discrepancies were observed. For example, *Streptococcus* species were underrepresented among MAGs, which is a pattern reported previously and attributed to high genomic similarity and strain-level diversity heterogeneity which complicate assembly and binning[6,7]. Additional differences arise from the use of distinct databases and taxonomic frameworks: MetaPhlAn relies on clade-specific marker genes, whereas FastANI and Mash use whole or draft genome comparisons against NCBI and RefSeq. Tool versioning may also contribute, as MetaPhlAn v3 was used here, while the more recent v4 includes an expanded reference genome and MAG catalogue.

Despite these differences, MAG-based analysis provided critical biological insights that cannot be obtained from read-based data alone. Genome-resolved analyses enabled species-level replication rate estimation and pangenome-based assessments of gene content dynamics, allowing us to interrogate how individual taxa grow, persist, and function over time. These analyses capture ecological strategies and population dynamics at both species and genus levels, providing mechanistic context for read-based observations and helping to distinguish active population expansion from passive persistence or decline.

Together, read-based and MAG-based results offer complementary perspectives on children oral microbiome development. Read-based profiles capture broad community restructuring across timepoints, whereas genome-resolved analyses reveal how specific populations respond to ecological pressures through changes in growth, functional capacity, and strain composition. The concordance between these layers, along with informative discrepancies, strengthens our inference of oral microbiome succession as a dynamic, species-specific process shaped by both ecological opportunity and host-associated factors across childhood.

**Functional profiling of oral microbiome**

Functional capacity was referred to the functional pathways inferred from microbial gene contents within the oral microbiome and was assessed using both reads-based and MAG-based approaches using multiple methods. For reads-based analysis, we focused on MetaCyc pathways which represent inferred metabolic capacity based on underlying gene contents and provide an interpretable framework for community-level functional comparisons. Specifically, stratified pathway profiles were used to assess overall functional shifts across timepoints which provided greater discriminatory power as confirmed in classification analyses compared to unstratified profiles. In contrast, unstratified functional profiles were used for ACE modelling to capture host genetic and environmental influences on overall community-level functional capacity rather than species-specific effects. We further characterised gene content variation in selected key species *Streptococcus mitis* using PanPhlAn, which reconstructs strain-level pangenomes directly from metagenomic reads by mapping them to a species-specific reference pangenome database. This enabled assessment of core and accessory gene presence–absence patterns and relative gene abundance across samples, providing insight into within-species functional heterogeneity and strain-level dynamics during oral microbiome development. Future work integrating additional functional layers such as KEGG and enzyme-based profiling may provide additional insights into the understanding of functional maturation in the oral microbiome.

In parallel, MAG-based functional annotation was performed using Anvi'o with the NCBI Clusters of Orthologous Groups (COGs) database to assign genes encoded in each MAG to broad functional categories, including metabolism, information processing, and cellular processes and signalling. By leveraging high-quality MAGs, anvi'o enabled genome-wide resolution of gene repertoire variation across strains. This facilitated gene cluster-based core and accessory analysis, allowing us to examine how functional potential is structured at the population level and how core and accessory functions are distributed among strains during childhood oral microbiome development. These analyses revealed species-specific patterns of gene gain, loss, and functional diversification that are not detectable from abundance-based or pathway-level profiling alone, further highlighting the dynamic and adaptive nature of oral microbial populations across childhood.

**MAGs within oral microbiome**

A key contribution of this study is the recovery of a large collection of high-quality oral MAGs, including genomes from poorly represented or previously uncultivated taxa such as members of the TM7 phylum. Given that only a small proportion of the oral microbiome is currently culturable, this expanded MAG catalogue represents a valuable resource that will enrich public databases and support future investigations into oral microbial ecology, evolution, and host interaction. Additionally, TM7 MAGs will contribute to future studies on whole genome assembly and characterisation of understudied lineages within this phylum.

MAG quality varied across developmental stages, with T1 assemblies showing slightly higher completeness, lower contamination, and longer contig N50 compared with T2–T3 (Extended Data Fig. 6). This pattern is expected with the finding of our study that early-life oral microbiomes are less complex, enabling more complete and contiguous genome reconstruction, whereas increasing microbial diversity and strain heterogeneity at later ages reduce assembly quality. These quality differences have implications for downstream analyses, as they reduce the number of high-quality MAGs available at T2 and T3. Measures such as replication rate and pangenome construction require high-quality genomes; therefore, we restricted these analyses to species with sufficient numbers of high-quality MAGs. Despite these constraints, our key conclusions remain robust. We applied consistent quality thresholds across timepoints and used comparable numbers of MAGs to ensure that observed patterns reflect true ecological progression rather than assembly difference. Additionally, randomisation procedures further confirmed the reliability of our results.

**ACE twin model on relative contributions of genetic and environmental factors**

The temporal fluctuations in A, C, and E components indicate that the relative contributions of genetic and environmental factors to the oral microbiome shift dynamically across childhood. These changes likely reflect developmental transitions in lifestyle, diet, behaviour, and oral hygiene practices but are not necessarily a continuous change across timepoints.

It is important to emphasise that the ACE model estimates the genetic (A), shared environmental (C), and unique environmental (E) components independently at each timepoint. ACE does not model a continuous developmental process; rather, it reflects the twin correlation structure based on the variance of microbial species and functions measured at each specific timepoint. Thus, differences across timepoints do not imply abrupt biological changes, but instead reflect how the relative contributions of genes and environment manifest at each temporal stage. Such example from our analysis is *Streptococcus parasanguinis*, where heritability is moderate at T1 and high at T3, yet appears negligible at T2. This pattern is statistically valid and biologically plausible. The toddler stage (T2, ~1.6 years) is a period of major environmental transitions with diversification of diet, establishment of tooth-brushing routines, daycare attendance, and strong parental structuring of daily behaviours. These factors can create a highly uniform shared environmental context across children, reducing detectable genetic variance and amplifying the C component. While the abundance of some species appears strongly shaped by environmental factors at T2, others remain more genetically influenced indicating species-specific features. Additional longitudinal sampling across more timepoints would help clarify fine-scale trajectories and transitions.

This classical ACE model has several assumptions. It assumes that all genetic effects are additive with no contribution from dominance or epistatic interactions and does not explicitly model gene–environment interaction or correlation. The ACE model infers genetic and environmental contributions indirectly from twin similarity patterns and does not resolve specific environmental exposures. The model relies on the equal environments assumption that monozygotic (MZ) and dizygotic (DZ) twins are assumed to experience shared environments to a similar extent and violation of this assumption such as enhanced microbial sharing among MZ twins could inflate genetic estimates. However, we found no evidence for such inflation in our analysis, and it remains informative for assessing the relative contributions of host genetics and environment, particularly when interpreted in a comparative and developmental context. At T1 and T2, MZ twins did not exhibit greater within-pair similarity than DZ twins, indicating minimal genetic influence in early life and arguing against such violation. Intraclass Correlation Coefficient (ICC) calculated directly from observed data prior to model fitting further showed stable MZ similarity across timepoints, rather than the increasing similarity expected if microbial sharing intensified with age (Supplementary Table 13). In contrast, within-pair similarity declined among DZ twins, highlighting an increasing contribution of host genetic factors later in childhood.

These patterns are consistent with maturation of the oral ecosystem, in which increasingly occupied and structured niches limit the establishment of newly encountered microbes. As oral epithelia and salivary environments mature, host-specific binding and immune factors increasingly constrain microbial persistence, promoting stable, individual-specific biofilm communities. Model robustness was further supported by AE versus ACE comparisons, which yielded consistent patterns across timepoints. Nevertheless, genetic effects estimated under the ACE framework should be interpreted as an upper bound. Future analyses will incorporate additional individual-level traits as covariates, including not only sex but also diet and oral hygiene, to further investigate their contributions to oral microbiome variation.

### References


1. Caufield, P. W. *et al.* Natural history of Streptococcus sanguinis in the oral cavity of infants: evidence for a discrete window of infectivity. *Infect Immun* 68, 4018–4023 (2000).
2. Mrowicka, M., Mrowicki, J., Dragan, G. & Majsterek, I. The importance of thiamine (vitamin B1) in humans. *Biosci Rep* 43, (2023).
3. Keshet, A. & Segal, E. Identification of gut microbiome features associated with host metabolic health in a large population-based cohort. *Nat Commun* 15, 9358 (2024).
4. Sheng, S. *et al.* Structural and Functional Alterations of Gut Microbiota in Males With Hyperuricemia and High Levels of Liver Enzymes. *Front Med (Lausanne)* 8, (2021).
5. Lamont, R. J., Koo, H. & Hajishengallis, G. The oral microbiota: dynamic communities and host interactions. *Nat Rev Microbiol* 16, 745–759 (2018).
6. L., E. J. *et al.* Supragingival Plaque Microbiome Ecology and Functional Potential in the Context of Health and Disease. *mBio* 9, 10.1128/mbio.01631-18 (2018).
7. Lefébure, T. & Stanhope, M. J. Evolution of the core and pan-genome of Streptococcus: positive selection, recombination, and genome composition. *Genome Biol* 8, R71 (2007).
